# A Systematic Review and Large-Scale tES and TMS Electric Field Modeling Study Reveals How Outcome Measure Selection Alters Results in a Person- and Montage-Specific Manner

**DOI:** 10.1101/2023.02.22.529540

**Authors:** Sybren Van Hoornweder, Marten Nuyts, Joana Frieske, Stefanie Verstraelen, Raf L.J. Meesen, Kevin A. Caulfield

## Abstract

**Background:** Electric field (E-field) modeling is a potent tool to examine the cortical effects of transcranial magnetic and electrical stimulation (TMS and tES, respectively) and to address the high variability in efficacy observed in the literature. However, outcome measures used to report E-field magnitude vary considerably and have not yet been compared in detail.

**Objectives:** The goal of this two-part study, encompassing a systematic review and modeling experiment, was to provide an overview of the different outcome measures used to report the magnitude of tES and TMS E-fields, and to conduct a direct comparison of these measures across different stimulation montages.

**Methods:** Three electronic databases were searched for tES and/or TMS studies reporting E-field magnitude. We extracted and discussed outcome measures in studies meeting the inclusion criteria. Additionally, outcome measures were compared via models of four common tES and two TMS modalities in 100 healthy younger adults.

**Results:** In the systematic review, we included 118 studies using 151 outcome measures related to E-field magnitude. Structural and spherical regions of interest (ROI) analyses and percentile-based whole-brain analyses were used most often. In the modeling analyses, we found that there was an average of only 6% overlap between ROI and percentile-based whole-brain analyses in the investigated volumes within the same person. The overlap between ROI and whole-brain percentiles was montage- and person-specific, with more focal montages such as 4×1 and APPS-tES, and figure-of-eight TMS showing up to 73%, 60%, and 52% overlap between ROI and percentile approaches respectively. However, even in these cases, 27% or more of the analyzed volume still differed between outcome measures in every analyses.

**Conclusions:** The choice of outcome measures meaningfully alters the interpretation of tES and TMS E-field models. Well-considered outcome measure selection is imperative for accurate interpretation of results, valid between-study comparisons, and depends on stimulation focality and study goals. We formulated four recommendations to increase the quality and rigor of E-field modeling outcome measures. With these data and recommendations, we hope to guide future studies towards informed outcome measure selection, and improve the comparability of studies.

## 1. Introduction

Electric field (E-field) modeling is a computational approach to estimate the amount of transcranial electrical stimulation (tES) and transcranial magnetic stimulation (TMS) that reaches the cortex [1, 2]. By segmenting an individual’s structural magnetic resonance imaging (MRI) scan into different tissue types such as skin, bone, cerebrospinal fluid (CSF), grey matter, and white matter, it is possible to simulate the magnitude of E-fields induced by tES and TMS. By doing so, E-field modeling provides a potent tool to individually examine the effects of noninvasive brain stimulation and to address the high variability in efficacy that is currently observed across individuals. In the past, E-field modeling has already helped researchers derive novel tES montages and identify dose-response relationships [3-8], has suggested optimal stimulation targets in clinical cohorts [9, 10], and has pinpointed which cortical regions are being stimulated by TMS [11]. In recent years, the introduction of software packages such as SimNIBS and ROAST has catalyzed the widespread use of E-field modeling [1, 2]. However, despite the standardization offered by these software packages, crucial experimental decisions made by researchers can vary considerably between studies, affecting study results [12-16]. In particular, an often-overlooked determinant of E-field modeling findings is the selected outcome measure to quantify E-field magnitude, which may affect interpretation.

To date, researchers have used numerous outcome measures to quantify the magnitude of E-fields produced by tES and TMS. For instance, E-field magnitude has been quantified both in a region of interest (ROI) and as a percentile of the total induced E-field magnitude. While both metrics pursue a common goal, they do so in a substantially different manner. Whereas the ROI focuses on a specific, user-defined brain region (e.g., the left primary motor cortex [M1]) [17-22], a percentile whole-brain approach studies E-fields across the entire brain and does not restrict the analyses to any region [23-25]. Moreover, even within the same approach, wide methodological variations prevail. Some studies have quantified E-field magnitude within spherical ROIs of considerably different radii, ranging between 0.5 and 45 mm, whereas others used different geometric shapes such as a cubical ROI [17-22].

While numerous studies have examined the importance of methodological factors such as MRI parameters [12, 14, 26, 27], head model detail [11, 27, 28] and meshing approach [2, 29], systematic investigation of the impact of modeling outcome measure choice on E-field magnitude results has not yet been achieved. Critically, the choice of outcome measure affects all modeling approaches even as more advanced segmentation and meshing techniques emerge, making it a key consideration both now and into the future. As focus shifts towards unraveling dose-response curves associated with E-field magnitude [30-34], it is crucial that we are comparing the same brain regions and volumes necessitating a critical evaluation of outcome measures.

Therefore, this two-part study set out to formally examine and compare the breadth and frequency of different outcome measures that have previously been used to quantify E-field magnitude. In Part 1, we conducted a systematic review of the literature to identify and describe all the different outcome measures that researchers have used to quantify tES and TMS induced E-field magnitude. While a recent narrative review reported the use of E-field modeling within the scope of a specific software package [35], we sought to systematically report which outcome measures have been used so far across all software packages and populations. In Part 2, to further facilitate the interpretation of previous work and inform future work, we used E-field modeling and examined the most common approaches extracted from the systematic review in a large open-source dataset of 100 healthy adults to elucidate the impact of outcome measure choice. As different modalities stimulate varying volumes of brain tissue and focus E-fields in different ways, we simulated and compared outcome measures across 4 tES and 2 TMS montages (600 total models).

## 2. Methods

### 2.1. Systematic Review: Eligibility, Search Strategy and Extracted Information

This review was conducted according to the Preferred Reporting Items for Systematic Reviews and Meta-Analyses (PRISMA) Statement [36]. We consulted three electronic databases (PubMed, Scopus and Web of Science) to examine how E-field magnitude is quantified. We included studies if they adhered to the following eligibility criteria: (1) full-text availability; (2) written in English; (3) modeling of tES and/or TMS in humans; (4) reporting E-field magnitude. Studies were excluded if insufficient information was provided to reproduce the E-field magnitude extraction procedure. Given the significant advances made in the modeling field in the past decade, we confined our literature search to 2012–2022, with the final search taking place on December 5^th^, 2022. Our search keys are shown in **Supplementary materials 1**. We extracted the approach used to quantify E-field magnitude from each included study. As we did not extract the outcome measures of the included studies, but rather the approach to obtain E-field magnitude, we did not perform risk of bias analyses.

### 2.2. Computational modeling

#### 2.2.1. Head Model Creation Overview

All simulations were performed in SimNIBS v4.0.0. To disentangle how different outcome measures affect E-field magnitude, we retrieved T1w and T2w structural MRI scans from 100 participants from the Human Connectome Project dataset (22–35 years, 50 females) [37]. Through the SimNIBS – Charm pipeline, these MRI scans were segmented and meshed into tetrahedral head models with 10 compartments (air, eyes, skin, muscle, compact bone, spongy bone, cerebrospinal fluid, veins, grey matter, and white matter; **Figure 1**) [1, 29]. All head models were visually inspected and we confirmed that there were no segmentation errors present.

**Figure 1.**
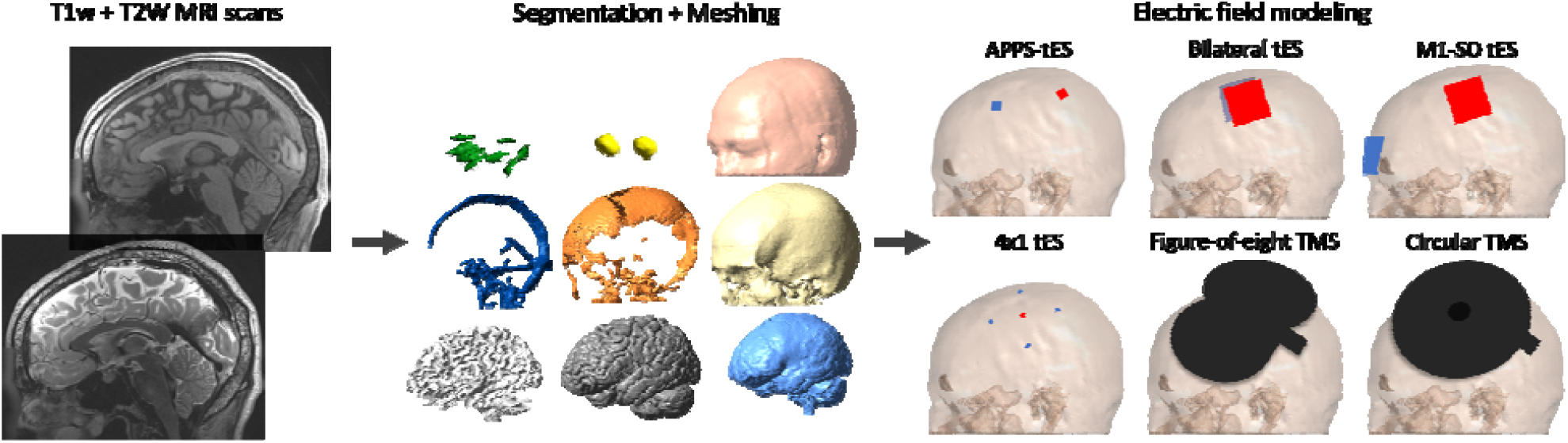
Electric field modeling pipeline. Per participant, we simulated four transcranial electrical stimulation (tES) and two transcranial magnetic stimulation (TMS) modalities.

#### 2.2.2. Electric Field Models

Four common tES and two common TMS montages targeting left M1 were modeled. Standard conductivity values were used (**Supplementary materials 2**). We performed all tES simulations at an intensity of 1 mA, and all TMS simulations at 50% stimulator output on a MagPro R30 machine (dI/dt = 75 × 10^6^ A/s). As the induced E-field magnitude is linearly proportional to the stimulation intensity, multiplying and dividing the obtained E-field strengths is a simple heuristic to convert the 1 mA and 50% stimulator output results to other intensities.

For tES, we simulated the following four montages (**Figure 1**). (1) APPS-tES consisting of two rectangular 1 by 1 cm electrodes, with the anode placed over CP3 and the cathode over FC3 [4]. (2) Bilateral tES consisting of two rectangular 5 by 5 cm electrodes, with the anode placed over C3 and the cathode over C4 [38]. (3) M1-SO tES consisting of two rectangular 5 by 5 cm electrodes, with the anode placed over C3 and the cathode over Fp2 [39]. (4) 4×1 tES consisting of 5 circular electrodes (r = 0.25 cm), with the anode placed over C3 and the four cathodes placed over FC3, C1, CP3 and C5 [3]. Here, the current going through each cathode was adjusted to -0.25 mA, to let 1 mA flow through the anode.

For TMS, we modeled two coils placed over C3 with a 45° angle to the midline (**Figure 1**). (1) Figure-of-eight TMS consisted of a MagVenture MC-B70 coil [40]. (2) Circular TMS consisted of a MagVenture MMC-140-II coil [40].

After a formal assessment (**Supplementary materials 3**), we replaced values exceeding the 99.9^th^ percentile E-field magnitude with the 99.9^th^ percentile E-field magnitude. This was done as higher values are prone to staircasing errors, and lower values underestimate the peak magnitude [41].

#### 2.2.3. Qualitative and Quantitative Assessments of Outcome Measures

Based on the systematic review (Cf., **3.1. Systematic Review Results**), we discussed the spherical, structural, and cubical ROI and whole-brain percentile approaches. Next to a qualitative discussion, all outcome measures except for the structural ROI were subjected to quantitative analyses. The structural ROI was not included in quantitative analyses as it is highly specific, based on the individual’s neuroanatomy, neurophysiology, and/or the atlas used to define it. Therefore, quantifying this method has limited applicability and generalizability. The spherical and cubical ROIs were centered at the subject-space transformed cortical projection of C3 [42].

For the spherical ROI, we examined how ROI size affects E-field magnitude by extracting the mean E-field magnitude obtained from ROIs with radii ranging between 0.5 and 45 mm. These radii were selected as outer limits based on our systematic review [21, 22]. We also extracted the percentile E-field magnitude from within these ROIs, with percentiles ranging from the 10^th^ to 99.9^th^ percentile. This range was selected to ensure a complete overview, with the 25^th^ and 100^th^ percentile being the lowest and highest percentiles used in the studies included in the systematic review [43, 44]. Finally, we correlated mean E-field magnitudes obtained from ROIs with differing sizes against each other per modality through Spearman’s correlations.

For the cubical ROI, we construed ROIs with identical volumes as the spherical ROIs to assess whether both approaches result in similar mean E-field magnitudes.

For the percentile-based whole-brain analyses, we extracted the 10^th^ to 99.9^th^ percentile E-field magnitude, the analyzed grey matter volume per percentile (mm^3^) and the Spearman’s correlation between different percentiles within the same montage.

Due to the computational nature of E-field models, no inferential statistical analyses were conducted.

### 2.3. Comparison of Different Outcome Measures

In addition to separate ROI and whole-brain percentile analyses, we directly compared common outcome measures per montage to aid interpretation of prior literature and guide future study methodologies. Based on the systematic review (Cf., **3.1. Systematic Review Results**), we compared the spherical ROI approach (radii ranging between 0.5 and 45 mm) against the percentile-based whole brain approach (percentiles ranging between the 10^th^ and 99.9^th^ percentile). In each modality and radius – percentile combination, we calculated the difference and Spearman’s correlation between E-field magnitudes obtained by both approaches, and the volumetric overlap. Volumetric overlap was quantified as the grey matter volume (mm^3^) included in both approaches, divided by the grey matter volume included by at least one approach.

## 3. Results

### 3.1. Systematic Review Results

We included 118 studies after removal of duplicates and title, abstract, and full-text screening. In these studies, E-field magnitude was reported 151 times as several studies reported more than one outcome measure (**Figure 2** and **Table 1**). Studies either extracted E-field magnitude from within a ROI (n=100, 71/29 tES/TMS) or the whole-brain (n=51, 16/35 tES/TMS).

**Table 1.**
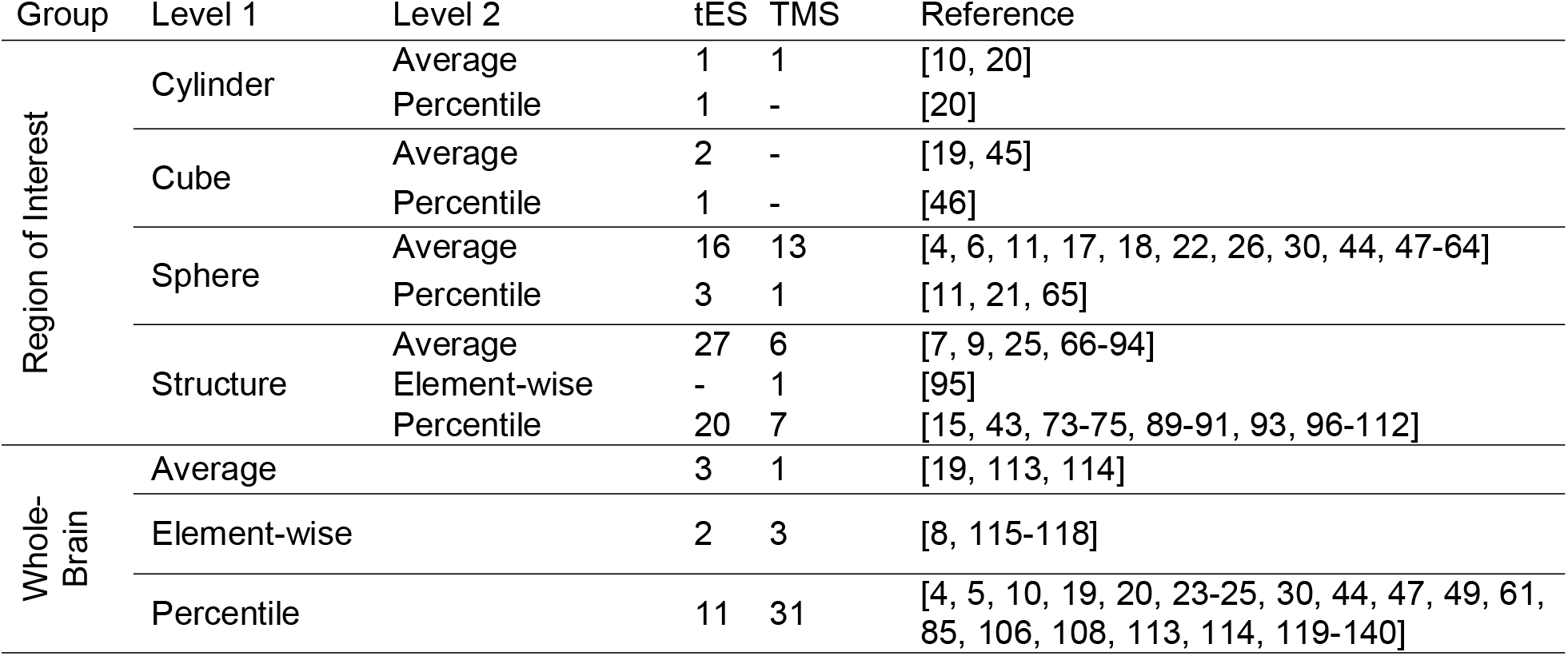
Electric field magnitude outcome measures

**Figure 2.**
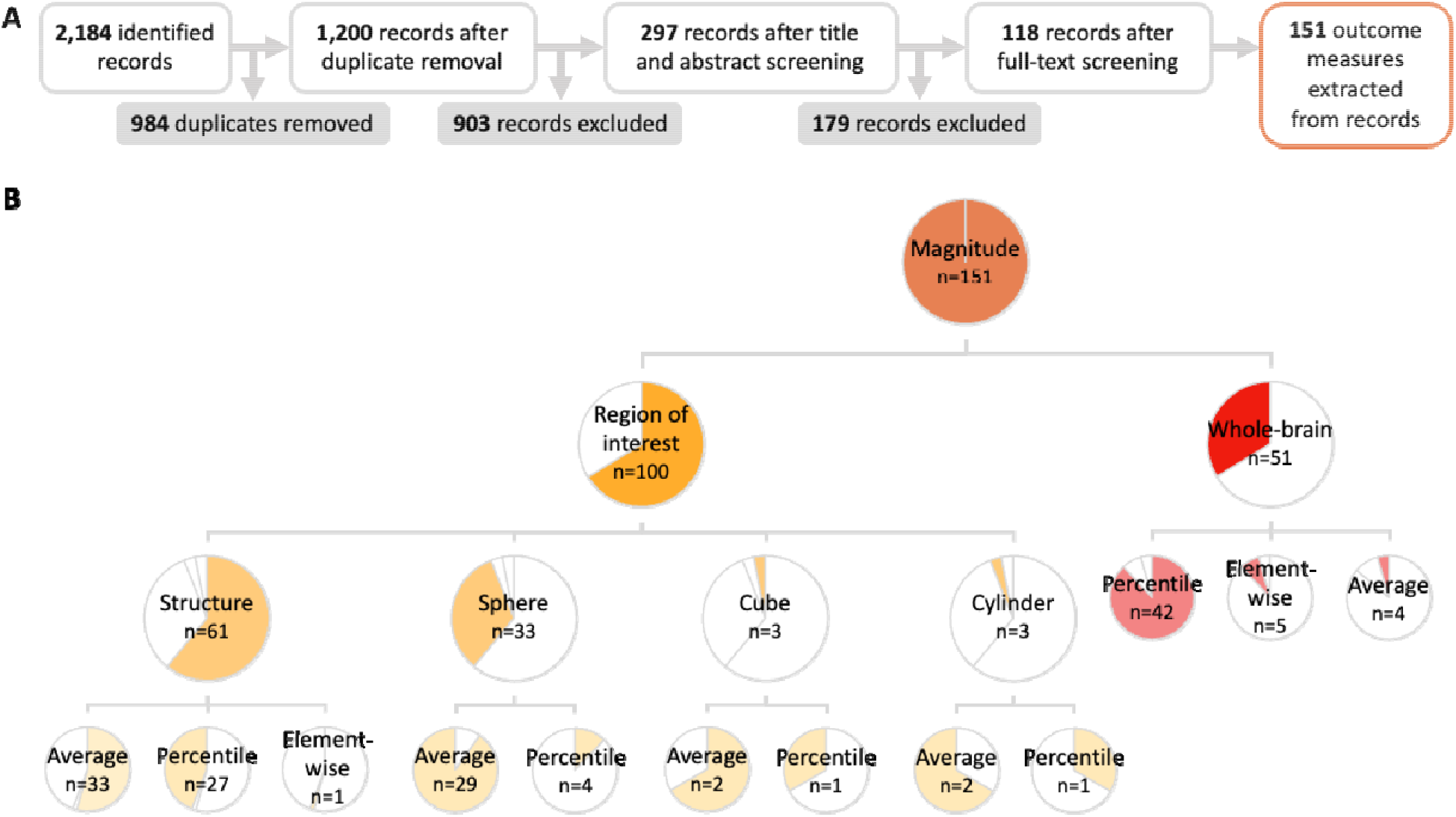
Extracted systematic review data. **A)** Flowchart of literature search and data screening. **B)** Tree diagram with pie charts displaying the frequency of each E-field magnitude outcome measure, relative to other outcome measures on the same level.

### 3.2. Region of Interest Outcome Measures Overview

ROI outcome measures involve extracting the E-field magnitude from within a ROI. First, we discuss spherical ROIs, as insights from these ROIs yield implications for the other ROIs.

#### 3.2.1. Spherical Region of Interest: Qualitative Results

In total, 33 studies extracted E-field magnitude via a spherical ROI (**Figure 2B** and **Table 1**). While most studies extracted the mean E-field magnitude from within the spherical ROI (n=29), a minority of studies extracted a percentile value from the spherical ROI (n=4).

A strength of spherical ROIs compared to structural ROIs and whole-brain approaches is that approximately the same area can be analyzed in terms of shape and volume across different brain regions, individuals, and studies. This facilitates a fairer head-to-head comparison of E-field magnitudes. Additionally, a spherical ROI is flexible in that its location can be derived from functional data (e.g., a hotspot obtained via TMS). However, a drawback is that spherical ROIs do not take cytoarchitectural data or complex neuroimaging data, beyond a single coordinate, into account. Also, while the spherical ROI may be identical in volume across individuals, the function and/or neurophysiology of the included brain volume can differ substantially. Likewise, defining the radius of a sphere is often arbitrary, leading to a wide variety in used radii, ranging from 0.5 to 45 mm in the included studies [21, 22]. Since each tES and TMS montage stimulates a different volume of grey matter tissue, the varying sizes of radii may or may not capture the actually stimulated network. Furthermore, as the grey matter tissue receiving the highest E-field magnitude can differ across persons within the same montage (cf.,**3.3.2. Percentile-Based Whole-Brain Outcome Measures: Quantitative Results**), spherical ROIs may capture the peak E-field magnitude in different degrees across different persons.

#### 3.3.2. Spherical Region of Interest: Quantitative Results

To assess how spherical ROI size impacts the obtained E-field magnitude, and, therefore, the interpretation of stimulation efficacy across tES and TMS protocols, we extracted the mean E-field magnitude and 10^th^ to 99.9^th^ percentile E-field magnitude in a percentile stepwise manner from a spherical ROI with radii ranging between 0.5 and 45 mm (**Figure 3**). Within each montage, we also assessed to what extent E-field magnitudes obtained by spherical ROIs with different sizes correlate to one another via Spearman’s correlations.

**Figure 3.**
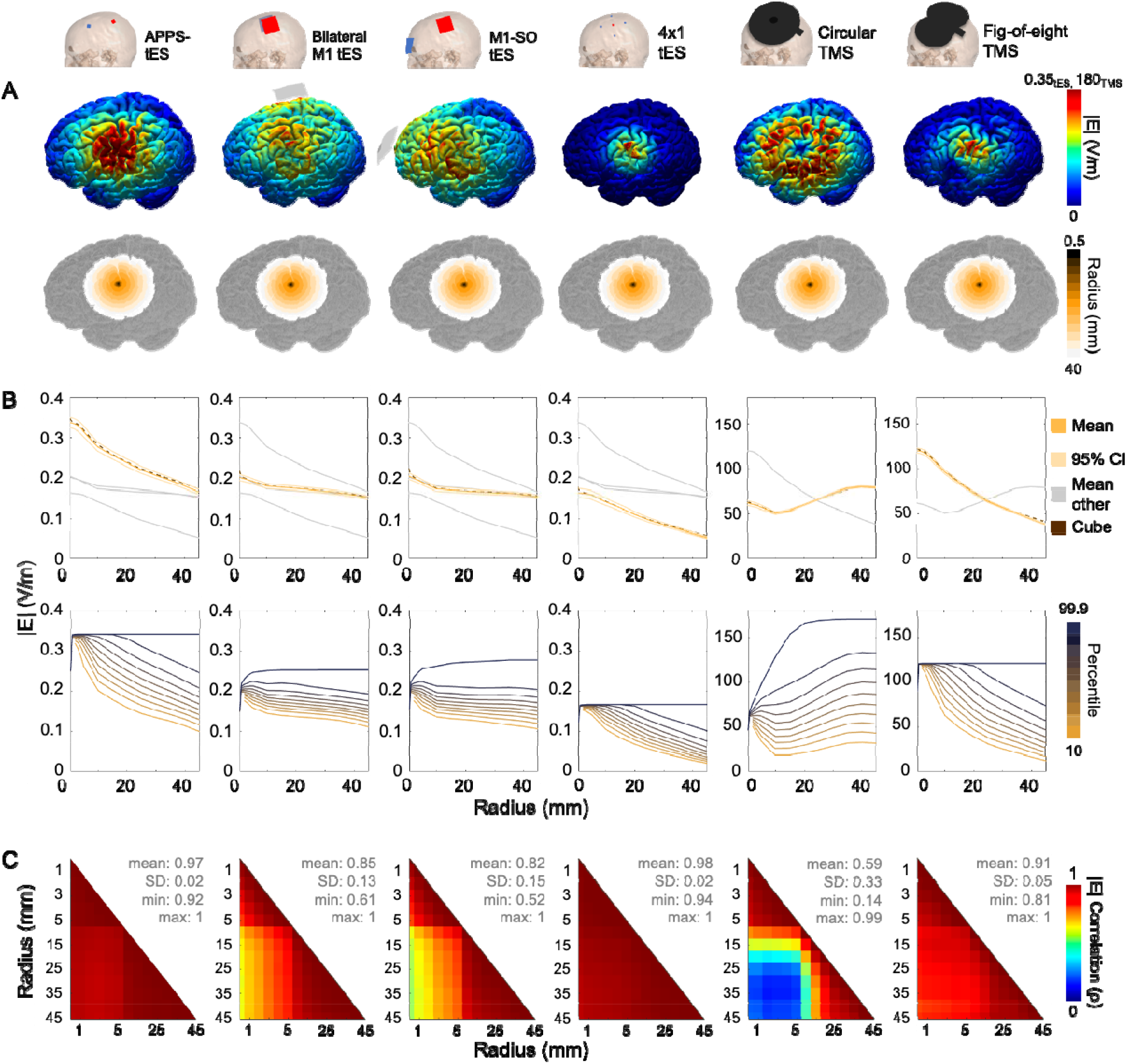
Effect of spherical region of interest (ROI) (radius in mm) size on E-field magnitude. **A)** Different montages with the induced E-field magnitude in an individual and the different spherical ROIs. **B**) Mean (upper row) and 10^th^ to 99.9^th^ percentile E-field magnitude (lower row) obtained by extracting the E-field in a sphere with radii ranging from 0.5 to 45 mm. In the upper row, we also show the mean E-field magnitude extracted by cubical ROIs with identical volumes to the spherical ROIs per radius size. **C**) Matrix of Spearman’s correlations between obtained mean E-field magnitudes across radii with varying sizes. SD = standard deviation.

ROI size affects the obtained E-field magnitude. This is crucial for future studies aiming to disentangle whether there is an optimal E-field magnitude to attain optimal behavioral and/or neurophysiological effects, as the answer strongly depends on the ROI size. An E-field magnitude obtained via a spherical ROI with a certain radius does not equal the same E-field magnitude obtained via a spherical ROI with a different radius. Therefore, the discussion must be limited to a particular ROI location and volume when considering the dose-response relationship between E-field magnitude and clinical effects.

For tES, larger ROIs resulted in lower E-field magnitudes (**Figure 3B**). Importantly, the effect of ROI size on E-field magnitude was montage-specific. The mean E-field magnitude obtained from more focal tES montages (i.e., APPS-tES and 4×1 tES) was most affected by the size of the ROI. Conversely, the peak E-field 99.9^th^ percentile remained stable in these montages across ROI size. For less focal tES montages, the obtained mean E-field magnitude depended less on ROI size. Here, the peak 99.9th percentile value became stable for spheres with radii exceeding 10 (bilateral M1 tES) and 20 mm (M1 – SO tES), suggesting that the peak E-field magnitude was not situated in the proximity of M1.

The differential susceptibility of more vs. less focal tES montages to ROI size can lead to interpretational pitfalls when comparing both tES classes. To illustrate, using a 5 mm radius sphere placed at the intended stimulation target of M1, APPS-tES induces considerably larger E-field magnitudes (mean ± SD = 0.32 ± 0.07 V/m), compared to bilateral M1 tES (0.19 ± 0.03 V/m). In contrast, when using a 45 mm radius ROI sphere, this difference is strongly attenuated (mean E-field magnitude APPS-tES: 0.17 ± 0.03 V/m, bilateral M1-tES: 0.15 ± 0.02 V/m). Thus, by varying the ROI size, one could either conclude that APPS-tES induces E-fields that are either 68% or 13% stronger than bilateral tES at the same stimulation intensity.

Reassuringly, within-montage correlation between E-field magnitudes extracted with ROIs with differing sizes was high, achieving mean Spearman’s ρ > 0.80 for all tES montages (**Figure 3C**). The lowest correlation was found for a spherical ROI with a 0.5 vs. 45 mm radius in the M1-SO montage (ρ = 0.52). This implies that, for within-subject designs, the impact of different ROI sizes seems negligible when using tES in most cases.

For TMS, the effect of ROI size on E-field magnitude depends on the coil type (**Figure 3B**). The slope of E-field magnitude induced by figure-of-eight TMS over radius resembles that of focal tES montages. For circular TMS, we observed an increase in mean E-field magnitude with ROI size, when ROI size exceeded 10 mm. This, combined with the observation that peak E-field values within the ROI also increase when larger ROIs are used, emphasizes that circular TMS does not produce maximal E-field magnitudes at the coil center, resulting in small ROIs not capturing the peak E-field magnitude.

The choice of sphere size affects the comparison of both TMS modalities. Using spheres with smaller radii (i.e., ≤ ~20 mm), one would conclude that figure-of-eight TMS induces higher E-field magnitudes. Using spheres with larger radii (i.e., ≥ ~30 mm), one would conclude that circular TMS induces higher magnitudes. Thus, similar to tES, whether there is a necessary or optimal E-field for clinical response is contingent on not only the stimulation parameters and location, but also on the specific outcome measure used.

The within-montage correlations of E-field magnitudes obtained across different spherical ROIs were high for figure-of-eight TMS (ρ > 0.81, for all correlations) (**Figure 3C**), implying that the choice of ROI size will not detrimentally affect between-subject comparisons for a given montage. However, correlations were weaker for circular TMS, where a minimal correlation coefficient of ρ = 0.14 was found for the correlation between E-field magnitude obtained by a 2 mm vs. 35 mm sphere. This implies that, even when measured within the same subject, there is barely a relationship between E-field magnitudes obtained by ROIs with different sizes.

Overall, these findings reveal that ROI size and mean-vs. percentile-based ROI methods are critical factors to consider in the E-field modeling domain. Particularly in the search of an optimal E-field strength, ROI size should be taken into account. Smaller ROIs placed at the intended stimulation target tend to result in larger mean E-field magnitudes, but often do not encapsulate the peak E-field magnitude as well as larger ROIs, particularly in non-focal montages. Lastly, aside from circular TMS, different ROI sizes are generally well-correlated to each other, which is reassuring for between-subject comparisons.

#### 3.2.3. Structural Region of Interest: Qualitative Results

In total, 61 studies extracted the E-field magnitude from a structure, defined as a geometrically irregular shape typically construed as an anatomical atlas or via neuroimaging data (e.g., Brodmann area or functional MRI activation map). The majority of these studies quantified E-field magnitude as the mean value (n=33) or a percentile value within the ROI (n=27).

The major advantage of structural ROIs is their flexibility towards specific research hypotheses or data. These ROIs allow to take the brain region’s cytoarchitecture into account via atlases such as the Human Connectome multi-modal Parcellation atlas, or can use neuroimaging data to define ROIs in a task-, subject- or group-specific manner [141]. Thus, structural ROIs can be tailored to specific research questions and data, whereas other ROIs may take different amounts of study-relevant vs. -irrelevant brain volumes into consideration across participants. When using structural ROIs, one should be aware that ROI size may influence the obtained E-field magnitude, in line with **Figure 3B**. However, when these ROIs are defined based on neuroimaging data, this is not necessarily a drawback.

Structural ROIs may also pose limitations beyond potentially overlooking other areas where relevant E-fields were generated. The uniqueness of structural ROIs in terms of size and shape, depending on the used data/atlas and the investigated brain region, undermines between-study and -region comparability. For instance, extracting E-field magnitudes via a structural ROI defined through a specific atlas will only provide insights with regard to that atlas and brain region. Therefore, the obtained E-field magnitude values are limited in terms of transferability to other studies, when different data or atlases are used. Moreover, as aforementioned in **2.2.3. Qualitative and Quantitative Assessments of Outcome Measures**, due to the geometric irregularity of structural ROIs, direct comparisons of different volumes, beyond what was achieved in **Figure 3**, are not meaningful due to their limited applicability.

#### 3.2.3. Cubical And Cylindrical Region of Interest Outcome Measures

Besides the structural and spherical ROIs, a minority of studies (n = 3 for both approaches) used cubical and cylindrical ROIs. While these methods resemble the spherical ROI in that they consist of constructing a geometrically regular shape around a given coordinate, they introduce an additional sensitivity to angular placement. Still, when combined with voxel-based neuroimaging, there can be compelling reasons to use a cubical ROI. For instance, Nandi et al. (2022) used a cubical ROI to extract E-field magnitude as this ROI matched the shape and size of the voxel they used for their magnetic resonance spectroscopy analyses [19].

As the cubical ROI can be useful, we compared this ROI to its spherical counterpart. We matched the volume of each spherical ROI (e.g., to match the 1 cm radius sphere, which has a volume of 4.189 cm^3^, we established a cube with a length, width and height of 1.612 cm) and extracted the mean E-field magnitude from the grey matter volume within our cubical ROIs, centered along the X, Y and Z axes. Our results (**Figure 3B**) indicate that when matched for volume, cubical and spherical ROIs result in close-to-identical mean E-field magnitudes. Thus, the same principles that we discussed for spherical ROIs should also apply to cubical ROIs.

### 3.3. Whole-Brain Outcome Measures Overview

In contrast to the regional specificity inherent to ROI analyses, whole-brain approaches remain agnostic to the spatial location receiving stimulation. In doing so, these analyses mitigate the risk of overlooking unexpected brain regions that receive relevant amounts of stimulation. This is a major strength assuming that the effects of tES and/or TMS are governed by a (monotonic) dose-response relationship in which a key factor is the peak E-field magnitude in any brain region.

#### 3.3.1. Percentile-Based Whole-Brain Outcome Measures: Qualitative Results

Most studies (n=42) used a percentile approach to quantify whole-brain E-field magnitude. This mitigates the risk of spatially poorly defining a ROI and overlooking the location that received the maximal E-field magnitude, potentially increasing the consistency of measured effects across participants and studies (**Figure 3B**). On the other hand, only extracting a percentile from E-field simulations introduces regional uncertainty. That is, the percentile approach is spatially undefined, resulting in it potentially providing information about unexpected regions that are not directly the stimulation target or providing information about different regions across participants.

#### 3.3.2. Percentile-Based Whole-Brain Outcome Measures: Quantitative Results

To disentangle how different percentile cut-offs affect the obtained E-field magnitude, we extracted the 10^th^ to 99.9^th^ percentile in a 10% incremental stepwise manner for all participants and montages. We also extracted and visualized the volume analyzed per percentile.

Although all approaches analyzed nearly identical volumes of grey matter tissue across the different percentiles (**Figure 4B**), visualization of the analyzed regions highlights that substantially different brain regions were analyzed by high percentile cut-offs across different tES and TMS modalities. For instance, **Figure 4A** indicates that the region receiving the highest E-field magnitude as a result of non-focal tES modalities is not located in M1. Perhaps even more concerning is the difference in analyzed regions across participants. **Figure 5** shows that the regions analyzed particularly by the 99.9^th^ percentile, can differ tremendously across persons, particularly in the non-focal tES montages. For instance, participant 88 (S88) shows peak E-fields in the proximity of M1 as a result of bilateral tES but not M1-SO tES, whereas participant 99 (S99) shows the opposite pattern. Thus, even the same outcome measure and threshold value can report E-field magnitudes in substantially different regions in a person-specific manner.

**Figure 4.**
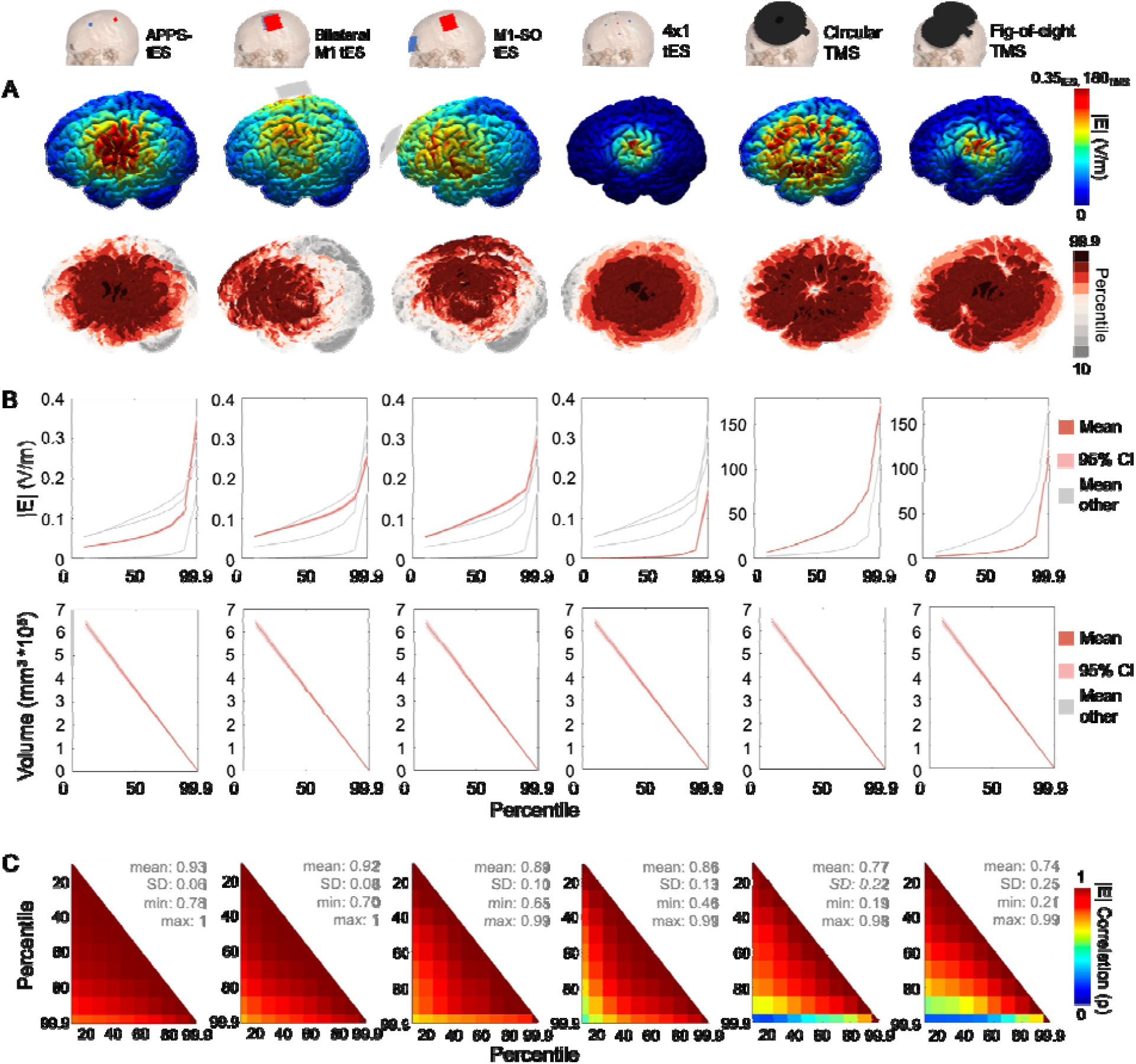
Effect of different percentiles on extracted E-field magnitude on the whole-brain level. **A)** Different montages with the induced E-field magnitude in an individual and the regions included in the different percentiles. **B**) 10^th^ to 99.9^th^ percentile E-field magnitude (upper row) and included volume (lower row). **C**) Matrix of Spearman’s correlations between obtained E-field magnitudes across varying percentiles.

**Figure 5.**
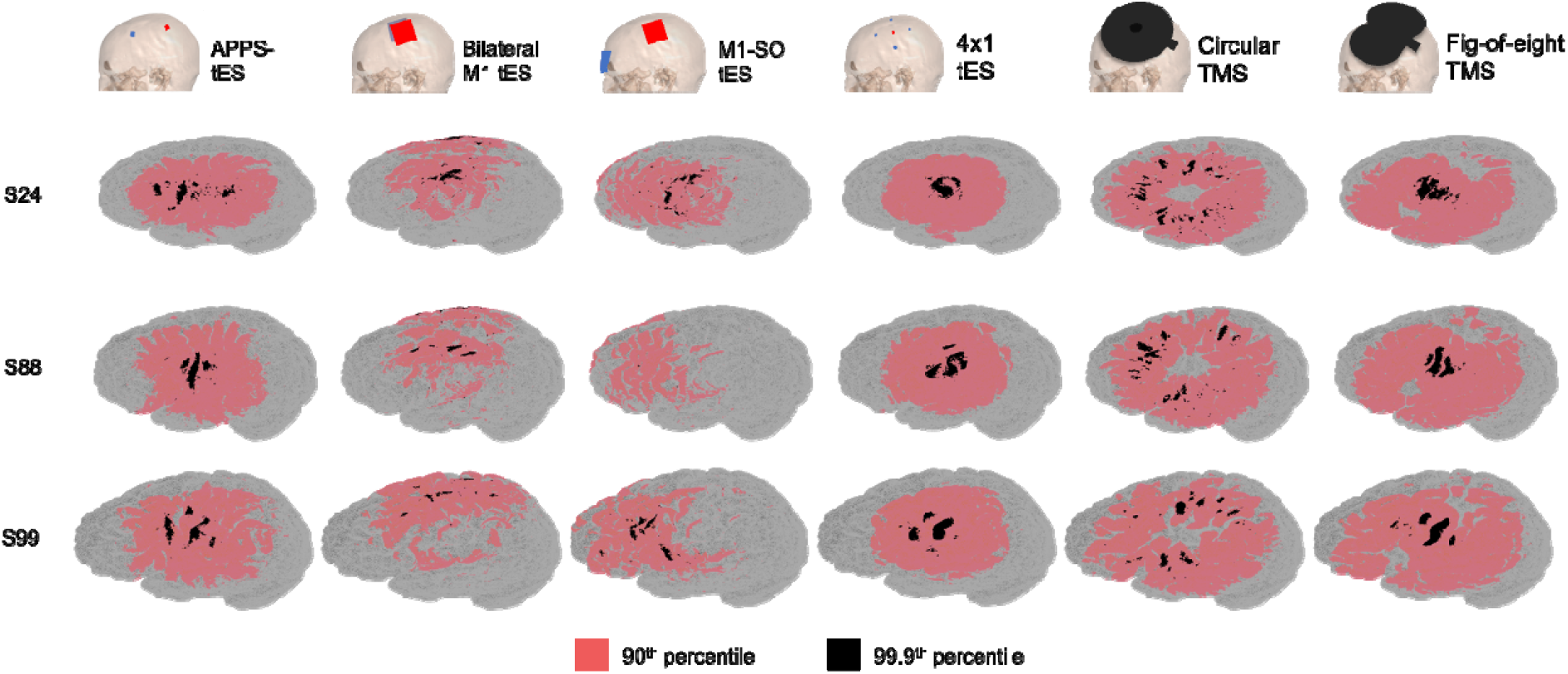
Volumes analyzed by the percentile-based outcome measure across all modalities in three subjects (S24, S88, and S99). Although the volumes analyzed by the 99.9^th^ percentile differ considerably for all modalities, this difference is most remarkable for bilateral and M1-SO tES.

For both TMS and tES, there was a steep increase in E-field magnitude from the 90^th^ to 99.9^th^ percentile (**Figure 4B**). E-field magnitude more than doubled for focal modalities, and increased by over 50% for non-focal modalities.

Overall for tES, the E-field magnitudes obtained by percentile approaches correlated strongly, with the mean correlation exceeding ρ = 0.86 for all modalities (**Figure 4C**). This is reassuring, as it implies that using different percentiles will likely not vastly impact conclusions of studies comparing between subjects. The lowest correlation value included the 99.9^th^ percentile E-field magnitude across all tES modalities, with 4×1 tES showing the lowest correlation (ρ = 0.46) between the 99.9^th^ and 10^th^ percentile. For TMS, the correlation between the 99.9^th^ percentile vs. all other percentiles is considerably lower than the correlation between the tES 99.9^th^ percentile vs. all other tES percentiles. Likely, this discrepancy stems from the fact that the magnetic field strength is more prone to decay with distance, while tES induced fields are governed by electro-conductive principles (**Figure 4B**).

#### 3.3.3. Element-Wise Whole Brain Outcome Measure

A minority of studies using a whole-brain approach extracted the element-wise E-field magnitude (n=5). While this method resembles percentile-based extraction, it is biased towards the spatial resolution of the head model (i.e., the number of tetrahedra in the case of SimNIBS). To illustrate, assuming that each grey matter element is identical in volume and the distribution of E-field magnitudes across elements is uniform, extracting the E-field magnitude of the 100^th^ highest element will result in a value similar to the 50^th^ percentile in a model with 200 elements (low spatial resolution) and a value similar to the 90^th^ percentile in a model with 1,000 elements (high spatial resolution).

#### 3.3.4. Mean Whole Brain Outcome Measure

Four studies extracted the mean E-field magnitude in the whole grey matter. Using this approach yields the disadvantage that, certainly for focal types of stimulation, it does not represent the E-field magnitude to which some regions were exposed.

### 3.4. Direct Comparison of Different Electric Field Magnitude Outcome Measures

To meaningfully inform future work, we directly compared two of the most common outcome measures: the mean spherical ROI approach and percentile-based whole-brain approach. To do so, we examined the difference and correlation between the obtained E-field magnitudes and the overlap between the analyzed brain volumes. For the spherical ROI, we used radii ranging between 0.5 and 45 mm. For the percentile-based whole-brain analyses, we used percentiles ranging between the 10^th^ and 99.9^th^ percentile.

First, we assessed to what extent the ROI and whole-brain approaches retain similar E-field magnitudes via Spearman’s correlations and E-field magnitude differences (%). While correlations inform how E-field magnitudes relate to the effects of stimulation across participants, E-field magnitude difference is a key consideration for research investigating absolute E-field magnitudes. Overall, correlations were highest for APPS-tES, where the mean correlation value was ρ = 0.86 (range = 0.76–0.98) and the lowest correlation value of ρ = 0.76 was obtained for the correlation between the 10^th^ percentile and a 4 mm spherical ROI. For bilateral M1-tES, the mean correlation value was ρ = 0.72 (range = 0.35–0.97). The highest correlations were present between the 90^th^ percentile and the larger spheres (i.e., spheres with radii ≥ 30 mm). With these comparisons, all correlations exceeded ρ = 0.95. For M1-SO tES, the mean Spearman’s value was 0.56 (range = 0.15–0.82); correlations were highest between the 80^th^ and 90^th^ percentile and spherical ROIs with radii exceeding 15 mm. Here, all correlation values exceeded ρ = 0.70. Notably, for M1-SO tES, the correlation between the 99.9^th^ percentile E-field magnitude and the spherical ROI was low-to-moderate, irrespective of ROI size (highest correlation: ρ = 0.54, lowest correlation: ρ = 0.15). Regarding 4×1 tES, the mean correlation value was ρ = 0.77 (0.42–0.99). The obtained correlation values resembled those of APPS-tES, although they were generally lower. Regarding circular TMS, the mean correlation value was ρ = 0.23 (range = -0.13–0.84). The highest correlations for circular TMS were present between spherical ROIs with radii ≥ 40 mm (lowest correlation: ρ = 0.83) and the 99^th^ percentile. Overall, circular TMS showed the weakest correlations between spherical E-field magnitudes and percentile-based E-field magnitudes, further corroborating that this type of TMS does not induce maximal E-field in the targeted ROI. For figure-of-eight TMS, the mean correlation value was ρ = 0.33 (range = -0.04–0.96). Here, a similar trend to the other more focal montages was present (i.e., high correlation values for the 99.9^th^ percentile against spherical ROIs of all sizes, and a gradual decrease in correlation values along with decreasing percentiles), although the obtained correlation values were notably lower for TMS vs. tES, pointing to a key consideration in future study designs depending on stimulation type.

Concerning E-field magnitude difference, the discrepancy was largest between E-field magnitudes in the 99.9^th^ percentile against all ROIs. Here, differences of up-to 338% were present (circular TMS, 99.9^th^ percentile vs. 10 mm ROI sphere). Although the differences with ROI-obtained E-field magnitudes were smaller for the other percentiles, the large overall difference between both approaches emphasizes the importance to only interpret and compare E-field magnitudes across studies using similar outcome measures.

The overlap of analyzed grey matter volumes is shown in **Figure 6, bottom row**. Overlap was generally low, with the mean overlap across all percentiles and ROI spheres not exceeding 6%. The highest overlap was present between the 90^th^ percentile and the large spherical ROIs with radii exceeding 30 mm. Here, an overlap between analyzed volumes of up to 60% (APPS-tES), 73% (4×1 tES) and 42% (Figure-of-eight TMS) was present. Of note, using ROIs of 10 mm yielded the greatest volumetric overlap with the 99.9^th^ percentiles, obtaining values of ≤ 21% overlap. **Figure 7**, which visualizes three spherical ROI volumes and two percentile whole-brain volumes further adds to this interpretation. Here, we show that the volume analyzed by the 90^th^ percentile indeed best resembles that of larger spherical ROIs. Moreover, based on the 99.9^th^ percentile, it becomes clear that the less focal types of stimulation induce their peak E-field magnitude outside of the intended stimulation target, while the spatially defined spherical ROI analyzes the E-field at the user-defined target.

**Figure 6.**
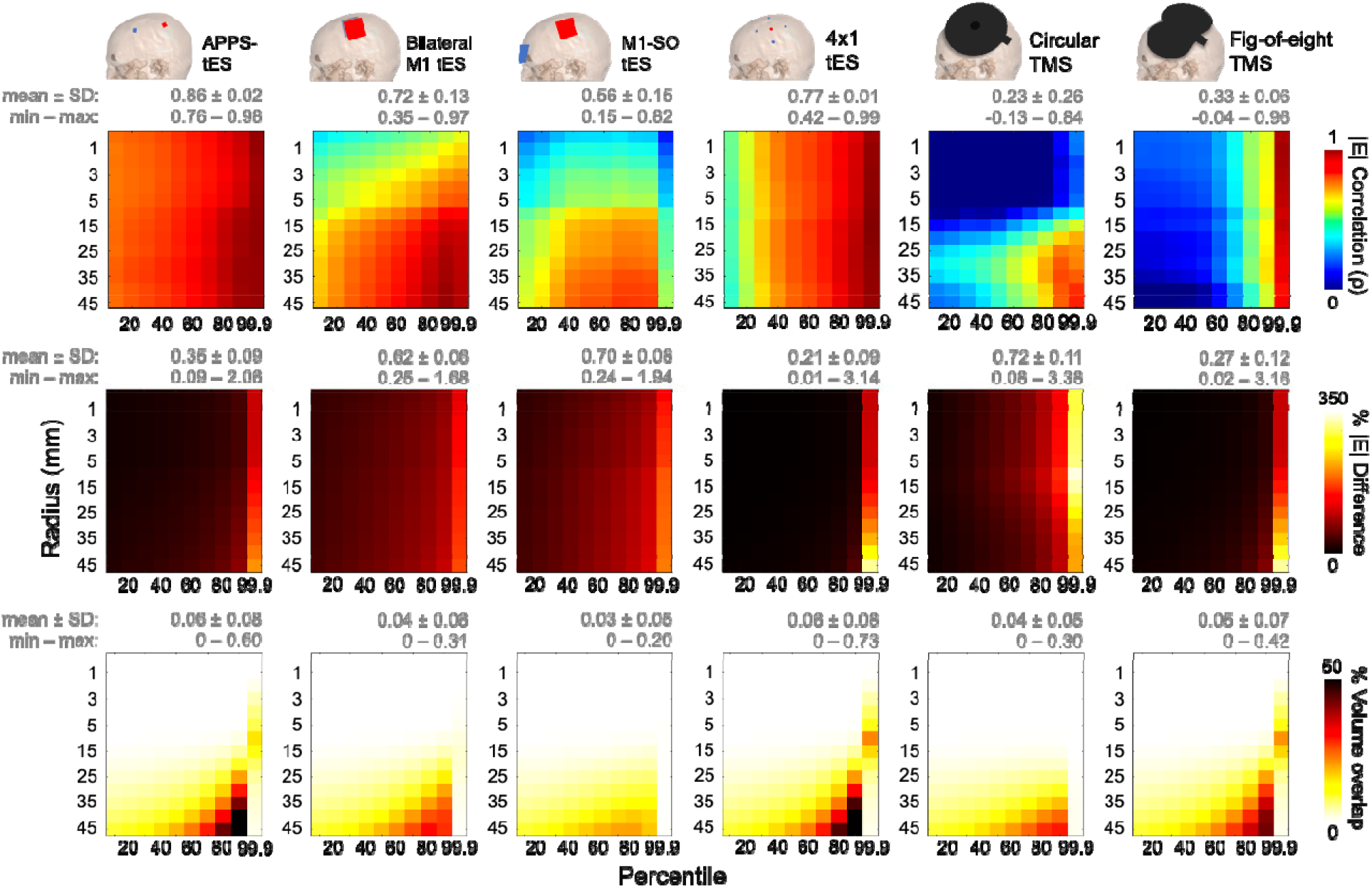
Comparison of spherical ROI and percentile-based whole brain E-field magnitude approaches in terms of correlation (upper row) and difference (middle row) in obtained E-field magnitudes and overlap in analyzed volume (bottom row).

**Figure 7.**
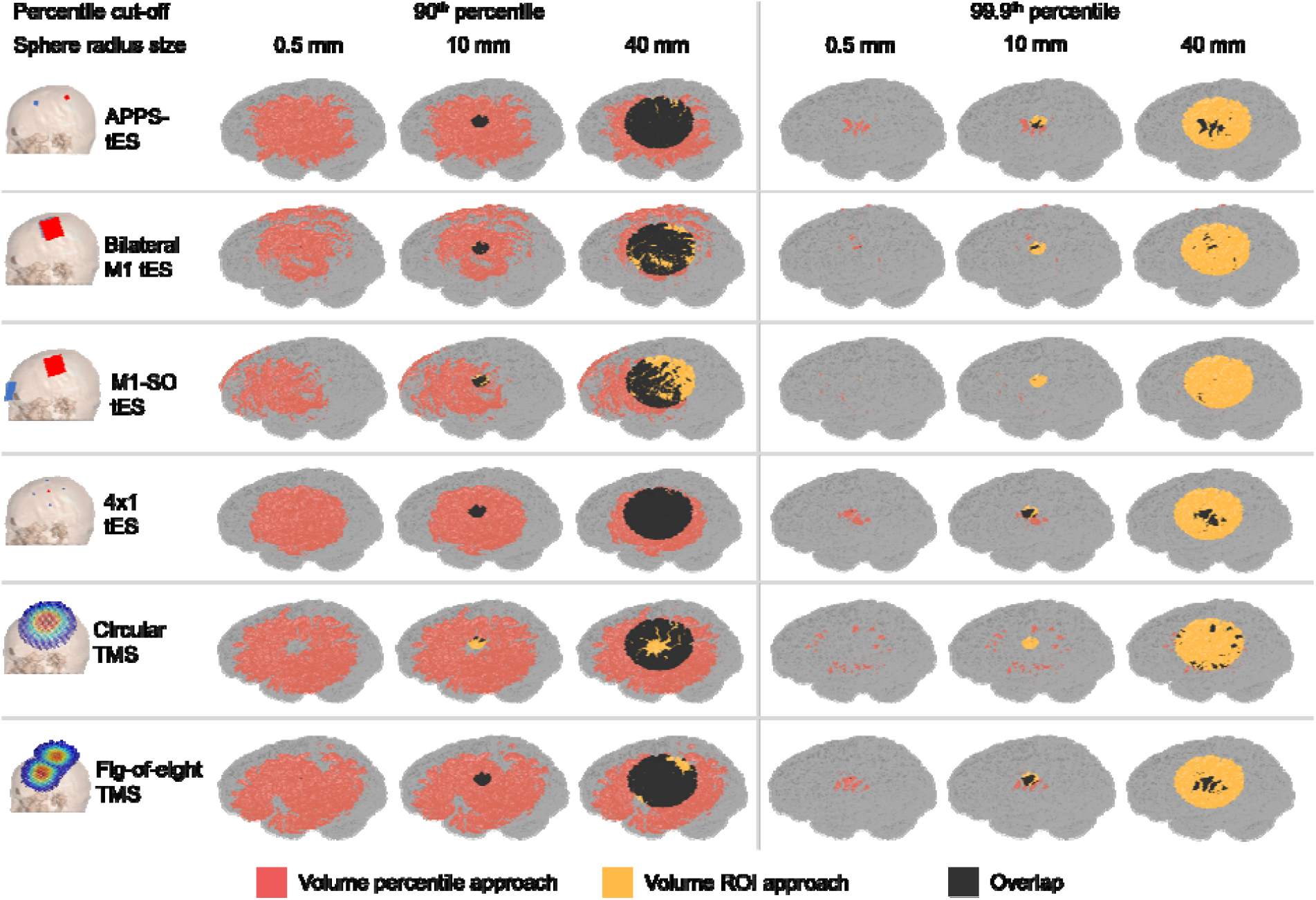
The volume analyzed by the spherical ROI approach (yellow), the percentile approach (red), and both approaches (i.e., the overlap) (black) in a representative participant.

## 4. Discussion

In this combined systematic review and E-field modeling study, we investigated how the selected outcome measure impacts E-field magnitude. In the 151 experiments reporting E-field modeling results that fit our systematic review criteria, outcome measures fell into two major categories: ROI (n = 100) and whole-brain approaches (n = 51). While most tES studies used the ROI approach (71% of studies), the whole-brain approach was used by the majority of TMS studies (68.6% of studies). We computed 600 E-field models in 100 healthy younger adults to further examine how E-field magnitude outcomes alter the interpretation of tES and TMS models that are more or less focal in stimulation diffusivity. Collectively, we computed 106,800 E-field magnitude related outcome measurements, finding that the selected outcome measure significantly impacts the interpretation of within- and between-subjects modeling. Specifically, for ROI-based outcomes, the wide range of sphere radii used in the literature examines volumes with a resolution level spanning from gyri to close-to-whole hemispheres. ROI size also interacts with stimulation focality; whereas the peak stimulation effects of focal forms of tES (APPS-tES and 4×1 tES) and TMS (figure-of-eight coil) are captured with focal ROIs placed at the stimulation target, the peak effects of less focal forms of tES (bilateral M1 and M1-SO) and TMS (circular coil) are better encapsulated with larger ROIs since these modalities do not always position the peak E-field at the intended target.

Regarding whole-brain approaches, we compared the E-field magnitude for each stimulation modality using percentiles ranging from the 10^th^ to the 99.9^th^ percentile in 10-percentile increments, finding high Spearman’s correlation values between each percentile with a mean value of at least ρ = 0.74 for all modalities. However, we found that percentile-based approaches analyzed different brain regions depending on the participant and the montage, introducing spatial uncertainty as a potential pitfall when interpreting the obtained E-field magnitudes. Namely, in less focal montages such as M1-SO tES, extracting E-field magnitudes through a high percentile threshold can result analyzing entirely different brain regions across persons. Therefore, it is important to consider how E-field outcome measure selection can not only alter group-level data but also data on the individual level.

The key question of whether there is an optimal E-field magnitude for dose-response relationships must be discussed within the context of each specific measure as we demonstrated wide variance of E-field modeling results based on the selected E-field measure. For instance, the optimal E-field for the 99.9^th^ percentile almost certainly differs from that at the 10^th^ percentile due to the volume examined and the consideration of different brain regions depending on the ROI size. Thus, more consistency and attention to E-field magnitude reporting measures is needed across studies.

We computed a percentile volumetric overlap between all 140 possible combinations per participant and modality to directly compare the mean ROI and percentile whole-brain approach. Strikingly, the mean overlap of percentile and ROI based outcome measures did not exceed 6%, suggesting that whole-brain percentile and spherical ROIs report the E-field magnitudes from fundamentally different brain regions. Overall, the low overlap underscores the critical importance of researchers selecting suitable E-field outcome measures to appropriately investigate their research questions. As we demonstrated here, selecting a suitable E-field outcome measure depends on the focality of the stimulation approach, the targeted region, and the volume of the targeted region or network. Crucially, these data directly comment on why it is important to be mindful when comparing across E-field modeling studies; even within the same person, stimulation parameters, and target, vastly different E-field magnitude findings can be extracted with mean ROI vs. percentile ROI vs. whole-brain percentile approaches. A valid critique is that the 6% overlap between the ROI and whole-brain percentile outcomes collapses across different volumes and therefore this overlap may be expected to be low. However, even when considering similar volumes between the ROI and whole-brain percentile approaches (e.g., the 90^th^ percentile and large spherical ROIs with radii exceeding 30 mm), the highest overlaps between outcome measures were still relatively low (60% for APPS-tES, 73% for 4×1 tES, and 42% for figure-of-eight TMS). Thus, even in the best case scenario, when comparing E-field magnitude results from ROI and whole-brain percentile approaches, analyzed volumes from at least 27% of the regions did not overlap.

### 4.1. Recommendations For Future Research

It is clear that the selected outcome measures can substantially affect the obtained E-field magnitude, and therefore, the interpretation of the results of computational simulations. The heterogeneity between outcome measures differs not only between ROI and whole brain approaches but also within measures such as the radius of a spherical ROI varying between

0.5–45 mm or the percentile cutoff varying between the 10^th^ to 99.9^th^ percentile. It is likely that these results are also translatable to the subcomponents of E-field magnitude (i.e., normal and tangential E-field strength), which use similar outcome measures.

Although ultimately, an outcome measure is always chosen in the context of specific research goals, there are some recommendations that can be made based on our review and data analyses that should be taken into account when deciding which outcome measure to use.

#### Recommendation #1: Use the Study Goal to Select the Most Suitable Outcome Measure

First, the choice for an outcome measure depends on the goal of the study and the available data. If neuroimaging data such as functional MRI is available, defining a structural ROI based on this data may be a viable approach. If the study’s goal is to compare E-field magnitudes in different neural regions or across persons, a spherical ROI may be best suited due to it analyzing nearly-identical grey matter volumes. It is important to weigh the pros and cons of the ROI and whole-brain percentile approaches as briefly summarized below and in **Table 2**.

**Table 2.**
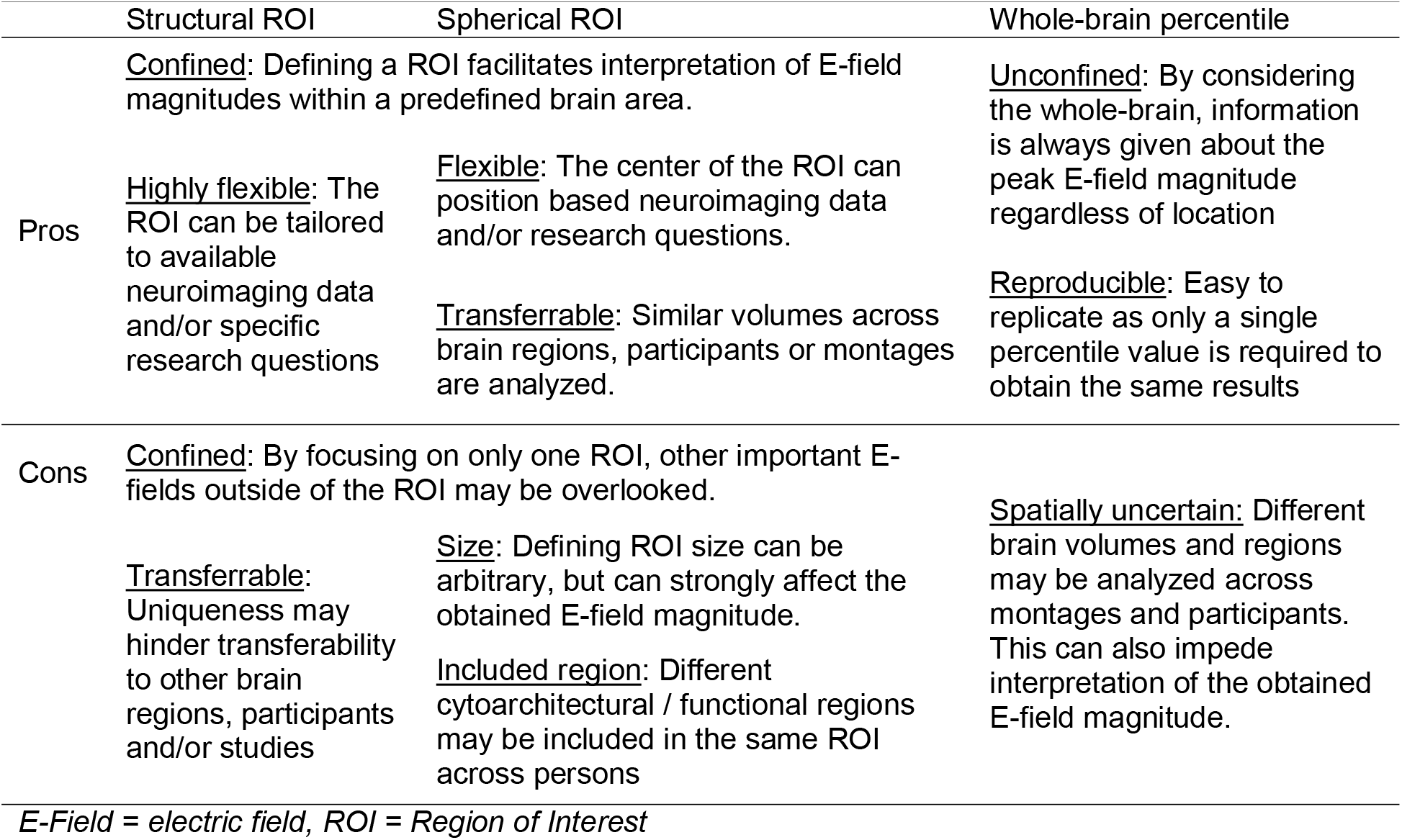
Pros and cons of the most common electric field modeling outcome measures

|  | Structural ROI | Spherical ROI | Whole-brain percentile |
| --- | --- | --- | --- |
| Pros | <u>Confined</u>: Defining a ROI facilitates interpretation of E-field magnitudes within a predefined brain area. |  | <u>Unconfined</u>: By considering the whole-brain, information is always given about the peak E-field magnitude regardless of location |
|  | <u>Highly flexible</u>: The ROI can be tailored to available neuroimaging data and/or specific research questions | <u>Flexible</u>: The center of the ROI can position based neuroimaging data and/or research questions. |  |
|  |  | <u>Transferrable</u>: Similar volumes across brain regions, participants or montages are analyzed. | <u>Reproducible</u>: Easy to replicate as only a single percentile value is required to obtain the same results |
| Cons | <u>Confined</u>: By focusing on only one ROI, other important E-fields outside of the ROI may be overlooked. |  |  |
|  | <u>Transferrable</u>: Uniqueness may hinder transferability to other brain regions, participants and/or studies | <u>Size</u>: Defining ROI size can be arbitrary, but can strongly affect the obtained E-field magnitude. | <u>Spatially uncertain</u>: Different brain volumes and regions may be analyzed across montages and participants. This can also impede interpretation of the obtained E-field magnitude. |
|  |  | <u>Included region</u>: Different cytoarchitectural / functional regions may be included in the same ROI across persons |  |

*E-Field = electric field, ROI = Region of Interest*

One characteristic that is inherent to all ROI outcome measures is the spatial confinement of these approaches. This is both a strength and weakness. It can be compelling when one is interested in a specific brain region. Conversely, this spatial confinement yields the danger that it is indifferent to the actual location that received maximal stimulation, which may not always coincide with the selected ROI but can play an important role to explain behavioral and/or neurophysiological effects. In addition, structural ROIs are highly unique which may impede between-region, -subject and/or -study comparisons and when using atlas-based structural ROIs, one may not be able to isolate the subregion of interest. On the other hand, spherical ROIs are biased towards the size of the sphere, particularly when the mean E-field magnitude is extracted, and may incorporate different volumes of task relevant grey matter across individuals.

Whole-brain percentile approaches may be best suited when researchers are interested in knowing the peak E-field magnitude irrespective of whether this coincides with the intended stimulation target. An advantage of whole-brain percentile outcome measures is that they produce identical outcomes by virtue of the mathematical threshold used, whereas some prior studies have used user-defined ROIs that could be prone to low inter-rater reliability. However, a limitation of the percentile approach is that it does not confirm target engagement of the intended stimulation region. Rather, with more diffuse stimulation methods such as M1-SO tES, the peak E-field is induced midway between electrodes. Thus, spherical ROI and whole-brain percentile approaches each have pros and cons and the correct choice depends on the researcher’s question.

#### Recommendation #2: Consider the Noninvasive Brain Stimulation Modality and Focality

Second, the optimal choice of outcome measure depends on which noninvasive brain stimulation modality is used. While ROI-based outcome measures are an excellent way to quantify the E-field magnitude in a specific region, their value is highest when analyzing more focal types of stimulation. In contrast, when less focal stimulation types are used, spatially confined ROIs placed directly underneath the electrodes or the coil center will often exclude the actual region that received the highest E-field magnitudes as the region of peak E-field intensity often differs from the target region and may differ across individuals.

#### Recommendation #3: The Dose-Response Relationship Between E-Field Magnitude and

*Clinical Outcome Must Only Be Compared Within a Singular Outcome Measure*

A primary finding of our systematic review is that researchers have used many different approaches to quantify E-field magnitude. With the powerful tool of E-field modeling, a question that many of us seek to answer is whether there is a positive dose-response relationship between the induced E-field magnitude and clinical response. It is enticing to consider these results monolithically and pursue an elegant statement such as, “The optimal E-field is V/m.” The simplicity of pursuing a singular value is appealing and would enable easier dissemination of individualized E-field dosing such as through dose-controlled tES, 2-Sample Prospective E-field Dosing (2-SPED), or applying an individualized TMS dose to induce a singular E-field at the cortical level across individuals [49, 51]. However, there are many reasons to believe that a singular optimal E-field value does not exist and that more nuance is necessary. For instance, a singular optimal E-field value would likely not apply in the same way across different brain regions due to varying neuronal populations, white matter tracts, and grey matter densities, among other variables [141-144]. Age, sex, diagnosis, and other typical demographical considerations likely also impact the optimal E-field dose [39, 64, 133, 145, 146]. Furthermore, the dynamic nature of the brain might further complicate things, as time-varying changes within a brain region of a single person may further modify the optimal E-field dose [147, 148]. Here, we added outcome measure as a key consideration to take into account in any discussion of optimal E-field dose. The E-field magnitudes extracted from the same models widely vary depending on the volume and regions considered. Thus, instead of the monolithic goal of a singular optimal E-field, we might instead work towards more nuanced goals taking many factors into consideration. In the future, we might come to the understanding that, “The optimal mean E-field for TMS, in a 5 mm radius spherical ROI centered over the motor hotspot as defined by TMS, in 50-to 70-year-old adult patients with ataxia, is V/m.” This value would almost certainly be different than an “optimal prefrontal E-field magnitude for TMS, measured by the 95^th^ percentile whole-brain approach, and in 20-to 40-year-old adult patients with depression.” Of course, additional refinement might further personalize our understanding of any “optimal” E-field value between individuals even when they have similar ages or diagnoses.

#### Recommendation #4: Time for Standardized E-Field Outcome Measure Reporting

Pursuing goals such as better understanding the dose-response relationship between E-field magnitude and therapeutic outcome necessitates that researchers report more standardized outcome measures between studies. As highlighted in *<u>Recommendation #3</u>* and throughout this study, comparing most ROI and whole-brain percentile measures is akin to comparing apples to oranges. Strikingly, it is even dissimilar to compare within some of these outcome measures across modalities, and even within the same participant and same E-field model when the ROI size has varied or the percentile has changed. Thus, in order to work toward more suitable comparisons across studies, it is important to consider how to improve the consistency of reporting across studies.

Notably, this recommendation does not conflict with *<u>Recommendation #1</u>* calling for researchers to choose the E-field outcome measure that best suits the experimental question. Rather, these reporting standards can apply to specific experimental questions which require different modeling outcome measures.

While deriving a comprehensive list of standard reporting procedures is beyond of the scope of this study and warrants a consensus-based approach, we propose a short list of preliminary guidelines to adopt across studies:

1. To state that a specific brain region was stimulated, researchers must use a ROI-based method and describe how the ROI was defined in the Methods section. We propose this guideline since the peak E-field intensity derived from the whole-brain percentile approach does not always coincide with the intended stimulation target, particularly with less focal forms of brain stimulation.
2. When defining a ROI, we recommend that researchers report the MNI coordinate that the ROI is centered on. In cases such as structural ROIs in which the researcher might individually define the ROI, an average MNI value should be provided when possible to aid in the reproducibility of findings and comparisons between studies.
3. Whether using a ROI or whole-brain percentile approach, the volume of the examined tissue should be reported. This recommendation seeks to allow for some degree of comparison between ROI and whole-brain percentile approaches since the reader should at least be able to determine whether a similar volume was analyzed. Researchers should ideally also visualize which regions are analyzed by a certain approach, with this being particularly important for percentile-based approaches to enable the reader to interpret the regions of the extracted volume.
4. Finally, we recommend all future studies to measure and report multiple outcome measures whenever possible. For instance, research applying M1-SO tES could use both the spherical ROI approach if one is interested in a specific region, and the percentile-based approach with complementary visualizations, to assess whether peak E-fields were induced in the intended region.

### 4.2. Conclusions

Outcome measures in the computational noninvasive brain modeling field have received minor attention in the past. Based on our systematic review and computational modeling of 106,800 E-field magnitude outcome measures across 100 participants and 600 tES and TMS E-field models, we show that different outcome measures substantially affect the obtained E-field magnitude and the analyzed brain region in a montage and person-specific manner. Therefore, one should only interpret and compare E-field magnitudes across studies when similar outcome measures are used. We formulated four recommendations (**Table 3**) for future research to ensure the informed selection of outcome measures. Our hope is that adopting these recommendations will enable future studies to avoid interpretational pitfalls and reduce the inconsistency of the used E-field outcome measures.

**Table 3.**
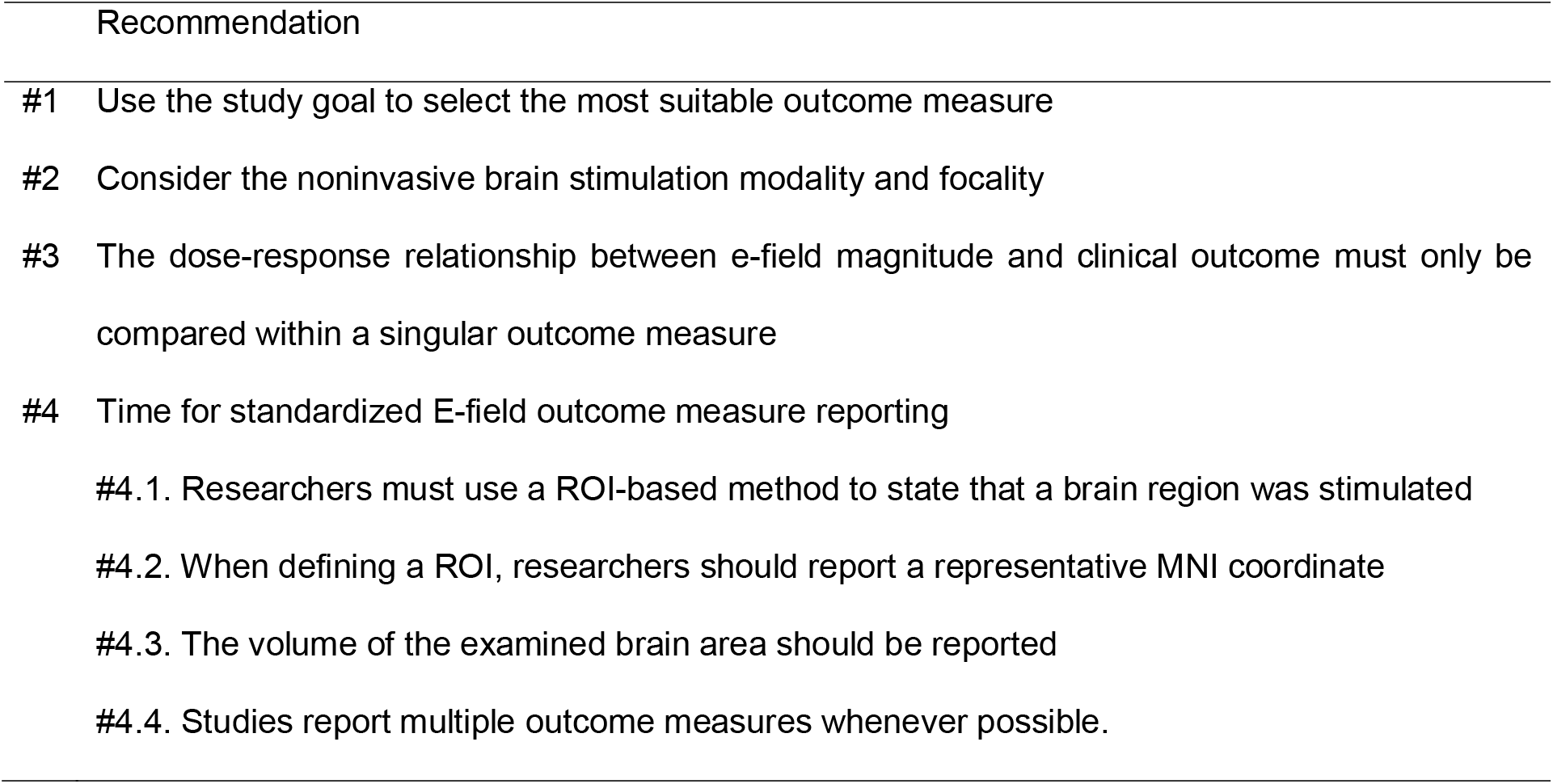
Overview of E-field modeling outcome measure related recommendations

## Funding

This work was supported by the Special Research Fund (BOF) of Hasselt University (BOF20KP18) (RM), Research Foundation Flanders (G1129923N) (SVH) an NIH NINDS F31 NRSA grant (1F31NS126019-01) (KAC). Data were provided by the Human Connectome Project, WU-Minn Consortium (Principal Investigators: David Van Essen and Kamil Ugurbil; 1U54MH091657) funded by the 16 NIH Institutes and Centers that support the NIH Blueprint for Neuroscience Research; and by the McDonnell Center for Systems Neuroscience at Washington University.

## Conflicts of interest

We confirm that all authors have no known conflicts of interest or competing interests associated with this publication and there has been no financial or personal relationship with other people / organizations that could inappropriately influence this work.

## Notes

### Competing Interest Statement

The authors have declared no competing interest.

## References

[1] Thielscher A, Antunes A, Saturnino GB. Field modeling for transcranial magnetic stimulation: A useful tool to understand the physiological effects of TMS? 2015 37th Annual International Conference of the IEEE Engineering in Medicine and Biology Society (EMBC). 2015:222–5.

[2] Huang Y, Datta A, Bikson M, Parra LC. Realistic volumetric-approach to simulate transcranial electric stimulation—ROAST—a fully automated open-source pipeline. J Neural Eng 2019;16(5):056006.

[3] Datta A, Bansal V, Diaz J, Patel J, Reato D, Bikson M. Gyri-precise head model of transcranial direct current stimulation: Improved spatial focality using a ring electrode versus conventional rectangular pad. Brain Stimul 2009;2(4):201–7.e1.

[4] Caulfield KA, George MS. Optimized APPS-tDCS electrode position, size, and distance doubles the on-target stimulation magnitude in 3000 electric field models. Sci Rep 2022;12(1):20116.

[5] Wischnewski M, Mantell KE, Opitz A. Identifying regions in prefrontal cortex related to working memory improvement: A novel meta-analytic method using electric field modeling. Neurosci Biobehav Rev 2021;130:147–61.

[6] Caulfield KA, Indahlastari A, Nissim NR, Lopez JW, Fleischmann HH, Woods AJ, et al. Electric Field Strength From Prefrontal Transcranial Direct Current Stimulation Determines Degree of Working Memory Response: A Potential Application of Reverse-Calculation Modeling? Neuromodulation 2020.

[7] Suen PJC, Doll S, Batistuzzo MC, Busatto G, Razza LB, Padberg F, et al. Association between tDCS computational modeling and clinical outcomes in depression: data from the ELECT-TDCS trial. Eur Arch Psychiatry Clin Neurosci 2021;271(1):101–10.

[8] Kasten FH, Duecker K, Maack MC, Meiser A, Herrmann CS. Integrating electric field modeling and neuroimaging to explain inter-individual variability of tACS effects. Nat Commun 2019;10(1):5427.

[9] Mantell KE, Sutter EN, Shirinpour S, Nemanich ST, Lench DH, Gillick BT, et al. Evaluating transcranial magnetic stimulation (TMS) induced electric fields in pediatric stroke. NeuroImage: Clinical 2021;29:102563.

[10] Minjoli S, Saturnino GB, Blicher JU, Stagg CJ, Siebner HR, Antunes A, et al. The impact of large structural brain changes in chronic stroke patients on the electric field caused by transcranial brain stimulation. NeuroImage: Clinical 2017;15:106–17.

[11] Weise K, Wartman WA, Knösche TR, Nummenmaa AR, Makarov SN. The effect of meninges on the electric fields in TES and TMS. Numerical modeling with adaptive mesh refinement. Brain Stimul 2022;15(3):654–63.

[12] Nielsen JD, Madsen KH, Puonti O, Siebner HR, Bauer C, Madsen CG, et al. Automatic skull segmentation from MR images for realistic volume conductor models of the head: Assessment of the state-of-the-art. Neuroimage 2018;174:587–98.

[13] Shahid S, Bikson M, Salman H, Wen P, Ahfock T. The value and cost of complexity in predictive modelling: Role of tissue anisotropic conductivity and fibre tracts in neuromodulation. J Neural Eng 2014;11:036002.

[14] Van Hoornweder S, Meesen RLJ, Caulfield KA. Accurate tissue segmentation from including both T1-weighted and T2-weighted MRI scans significantly affect electric field simulations of prefrontal but not motor TMS. Brain Stimulation: Basic, Translational, and Clinical Research in Neuromodulation 2022;15(4):942–5.

[15] Mikkonen M, Laakso I. Effects of posture on electric fields of non-invasive brain stimulation. Phys Med Biol 2019;64(6):065019.

[16] Opitz A, Yeagle E, Thielscher A, Schroeder C, Mehta AD, Milham MP. On the importance of precise electrode placement for targeted transcranial electric stimulation. Neuroimage 2018;181:560–7.

[17] Saturnino GB, Madsen KH, Thielscher A. Optimizing the electric field strength in multiple targets for multichannel transcranial electric stimulation. J Neural Eng 2021;18(1).

[18] Antonenko D, Thielscher A, Saturnino GB, Aydin S, Ittermann B, Grittner U, et al. Towards precise brain stimulation: Is electric field simulation related to neuromodulation? Brain Stimul 2019;12(5):1159–68.

[19] Nandi T, Puonti O, Clarke W, Nettekoven C, Barron H, Kolasinski J, et al. tDCS induced GABA change is associated with the simulated electric field in M1, an effect mediated by grey matter volume in the MRS voxel. Brain Stimul 2022.

[20] Rampersad SM, Janssen AM, Lucka F, Ü A, Lanfer B, Lew S, et al. Simulating Transcranial Direct Current Stimulation With a Detailed Anisotropic Human Head Model. IEEE Trans Neural Syst Rehabil Eng 2014;22(3):441–52.

[21] Saturnino GB, Antunes A, Thielscher A. On the importance of electrode parameters for shaping electric field patterns generated by tDCS. Neuroimage 2015;120:25–35.

[22] Zhang H, Gomez LJ, Guilleminot J. Uncertainty quantification of TMS simulations considering MRI segmentation errors. J Neural Eng 2022;19(2):026022.

[23] Laakso I, Mikkonen M, Koyama S, Hirata A, Tanaka S. Can electric fields explain inter-individual variability in transcranial direct current stimulation of the motor cortex? Sci Rep 2019;9(1):626.

[24] Saturnino GB, Madsen KH, Siebner HR, Thielscher A. How to target inter-regional phase synchronization with dual-site Transcranial Alternating Current Stimulation. Neuroimage 2017;163:68–80.

[25] Antonenko D, Grittner U, Puonti O, Flöel A, Thielscher A. Estimation of individually induced e-field strength during transcranial electric stimulation using the head circumference. Brain Stimul 2021;14(5):1055–8.

[26] Van Hoornweder S, Meesen R, Caulfield KA. On the importance of using both T1-weighted and T2-weighted structural magnetic resonance imaging scans to model electric fields induced by non-invasive brain stimulation in SimNIBS. Brain Stimul 2022.

[27] Huang Y, Liu AA, Lafon B, Friedman D, Dayan M, Wang X, et al. Measurements and models of electric fields in the in vivo human brain during transcranial electric stimulation. Elife 2017;6:e18834.

[28] Khadka N, Bikson M. Role of skin tissue layers and ultra-structure in transcutaneous electrical stimulation including tDCS. Phys Med Biol 2020;65(22):225018.

[29] Puonti O, Van Leemput K, Saturnino GB, Siebner HR, Madsen KH, Thielscher A. Accurate and robust whole-head segmentation from magnetic resonance images for individualized head modeling. Neuroimage 2020;219:117044.

[30] Turi Z, Hananeia N, Shirinpour S, Opitz A, Jedlicka P, Vlachos A. Dosing Transcranial Magnetic Stimulation of the Primary Motor and Dorsolateral Prefrontal Cortices With Multi-Scale Modeling. Front Neurosci 2022;16.

[31] Alekseichuk I, Wischnewski M, Opitz A. A minimum effective dose for (transcranial) alternating current stimulation. Brain Stimul 2022;15(5):1221–2.

[32] Wischnewski M, Mantell KE, Opitz A. Identifying regions in prefrontal cortex related to working memory improvement: A novel meta-analytic method using electric field modeling. Neuroscience and biobehavioral reviews 2021;130:147–61.

[33] Jamil A, Batsikadze G, Kuo HI, Labruna L, Hasan A, Paulus W, et al. Systematic evaluation of the impact of stimulation intensity on neuroplastic after-effects induced by transcranial direct current stimulation. The Journal of physiology 2017;595(4):1273–88.

[34] Chhatbar PY, Ramakrishnan V, Kautz S, George MS, Adams RJ, Feng W. Transcranial direct current stimulation post-stroke upper extremity motor recovery studies exhibit a dose– response relationship. Brain Stim 2016;9(1):16–26.

[35] Nasimova M, Huang Y. Applications of open-source software ROAST in clinical studies: A review. Brain Stimulation: Basic, Translational, and Clinical Research in Neuromodulation 2022;15(4):1002–10.

[36] Moher D, Liberati A, Tetzlaff J, Altman DG. Preferred reporting items for systematic reviews and meta-analyses: the PRISMA statement. PLoS Med 2009;6(7):e1000097.

[37] Van Essen DC, Ugurbil K, Auerbach E, Barch D, Behrens TE, Bucholz R, et al. The Human Connectome Project: a data acquisition perspective. Neuroimage 2012;62(4):2222–31.

[38] Nissim NR, O’Shea A, Indahlastari A, Telles R, Richards L, Porges E, et al. Effects of in-Scanner Bilateral Frontal tDCS on Functional Connectivity of the Working Memory Network in Older Adults. Front Aging Neurosci 2019;11.

[39] Ghasemian-Shirvan E, Mosayebi Samani M, Farnad L, Kuo M-F, Meesen R, Nitsche M. Age-dependent non-linear neuroplastic effects of cathodal tDCS in the elderly population; a titration study. Brain Stimul 2022;15.

[40] Drakaki M, Mathiesen C, Siebner HR, Madsen K, Thielscher A. Database of 25 validated coil models for electric field simulations for TMS. Brain Stimulation: Basic, Translational, and Clinical Research in Neuromodulation 2022;15(3):697–706.

[41] Soldati M, Laakso I. Effect of Electrical Conductivity Uncertainty in the Assessment of the Electric Fields Induced in the Brain by Exposure to Uniform Magnetic Fields at 50 Hz. IEEE Access 2020;8:222297–309.

[42] Okamoto M, Dan H, Sakamoto K, Takeo K, Shimizu K, Kohno S, et al. Three-dimensional probabilistic anatomical cranio-cerebral correlation via the international 10-20 system oriented for transcranial functional brain mapping. Neuroimage 2004;21(1):99–111.

[43] Fiocchi S, Chiaramello E, Marrella A, Bonato M, Parazzini M, Ravazzani P. Modelling of magnetoelectric nanoparticles for non-invasive brain stimulation: a computational study. J Neural Eng 2022;19(5):056020.

[44] Ciechanski P, Carlson HL, Yu SS, Kirton A. Modeling Transcranial Direct-Current Stimulation-Induced Electric Fields in Children and Adults. Front Hum Neurosci 2018;12:268.

[45] Caulfield KA, Badran BW, DeVries WH, Summers PM, Kofmehl E, Li X, et al. Transcranial electrical stimulation motor threshold can estimate individualized tDCS dosage from reverse-calculation electric-field modeling. Brain Stimul 2020;13(4):961–9.

[46] Parazzini M, Fiocchi S, Cancelli A, Cottone C, Liorni I, Ravazzani P, et al. A Computational Model of the Electric Field Distribution due to Regional Personalized or Nonpersonalized Electrodes to Select Transcranial Electric Stimulation Target. IEEE Trans Biomed Eng 2017;64(1):184–95.

[47] Marquardt L, Kusztrits I, Craven AR, Hugdahl K, Specht K, Hirnstein M. A multimodal study of the effects of tDCS on dorsolateral prefrontal and temporo-parietal areas during dichotic listening. Eur J Neurosci 2021;53(2):449–59.

[48] Caulfield KA, Li X, George MS. A reexamination of motor and prefrontal TMS in tobacco use disorder: Time for personalized dosing based on electric field modeling? Clin Neurophysiol 2021;132(9):2199–207.

[49] Van Hoornweder S A Caulfield K, Nitsche M, Thielscher A L J Meesen R. Addressing transcranial electrical stimulation variability through prospective individualized dosing of electric field strength in 300 participants across two samples: the 2-SPED approach. J Neural Eng 2022;19(5):056045.

[50] Caulfield KA, Badran BW, Li X, Bikson M, George MS. Can transcranial electrical stimulation motor threshold estimate individualized tDCS doses over the prefrontal cortex? Evidence from reverse-calculation electric field modeling. Brain Stimulation: Basic, Translational, and Clinical Research in Neuromodulation 2020;13(4):1150–2.

[51] Evans C, Bachmann C, Lee JSA, Gregoriou E, Ward N, Bestmann S. Dose-controlled tDCS reduces electric field intensity variability at a cortical target site. Brain Stimul 2020;13(1):125–36.

[52] Leunissen I, Van Steenkiste M, Heise K-F, Monteiro TS, Dunovan K, Mantini D, et al. Effects of beta-band and gamma-band rhythmic stimulation on motor inhibition. iScience 2022;25(5).

[53] Klaus J, Schutter DJLG. Electrode montage-dependent intracranial variability in electric fields induced by cerebellar transcranial direct current stimulation. Sci Rep 2021;11(1):22183.

[54] Gomez LJ, Dannhauer M, Peterchev AV. Fast computational optimization of TMS coil placement for individualized electric field targeting. Neuroimage 2021;228:117696.

[55] Soleimani G, Kupliki R, Bodurka J, Paulus MP, Ekhtiari H. How structural and functional MRI can inform dual-site tACS parameters: A case study in a clinical population and its pragmatic implications. Brain Stimul 2022;15(2):337–51.

[56] Lohse A, Meder D, Nielsen S, Lund AE, Herz DM, Løkkegaard A, et al. Low-frequency transcranial stimulation of pre-supplementary motor area alleviates levodopa-induced dyskinesia in Parkinson’s disease: a randomized cross-over trial. Brain Commun 2020;2(2):fcaa147.

[57] Caulfield KA, Fleischmann HH, Cox CE, Wolf JP, George MS, McTeague LM. Neuronavigation maximizes accuracy and precision in TMS positioning: Evidence from 11,230 distance, angle, and electric field modeling measurements. Brain Stimul 2022;15(5):1192–205.

[58] Zhang BBB, Stöhrmann P, Godbersen GM, Unterholzner J, Kasper S, Kranz GS, et al. Normal component of TMS-induced electric field is correlated with depressive symptom relief in treatment-resistant depression. Brain Stimul 2022;15(5):1318–20.

[59] Liu ML, Karabanov AN, Piek M, Petersen ET, Thielscher A, Siebner HR. Short periods of bipolar anodal TDCS induce no instantaneous dose-dependent increase in cerebral blood flow in the targeted human motor cortex. Sci Rep 2022;12(1):9580.

[60] Dmochowski JP, Datta A, Huang Y, Richardson JD, Bikson M, Fridriksson J, et al. Targeted transcranial direct current stimulation for rehabilitation after stroke. Neuroimage 2013;75:12–9.

[61] Janssen AM, Oostendorp TF, Stegeman DF. The coil orientation dependency of the electric field induced by TMS for M1 and other brain areas. J Neuroeng Rehabil 2015;12(1):47.

[62] Janssen AM, Oostendorp TF, Stegeman DF. The effect of local anatomy on the electric field induced by TMS: evaluation at 14 different target sites. Med Biol Eng Comput 2014;52(10):873–83.

[63] Mosayebi-Samani M, Jamil A, Salvador R, Ruffini G, Haueisen J, Nitsche MA. The impact of individual electrical fields and anatomical factors on the neurophysiological outcomes of tDCS: A TMS-MEP and MRI study. Brain Stimul 2021;14(2):316–26.

[64] Van Hoornweder S, Debeuf R, Verstraelen S, Meesen R, Cuypers K. Unravelling Ipsilateral Interactions Between Left Dorsal Premotor and Primary Motor Cortex: A Proof of Concept Study. Neuroscience 2021;466:36–46.

[65] Jog M, Anderson C, Kim E, Garrett A, Kubicki A, Gonzalez S, et al. A novel technique for accurate electrode placement over cortical targets for transcranial electrical stimulation (tES) clinical trials. J Neural Eng 2021;18(5).

[66] Rezaee Z, Ruszala B, Dutta A. A computational pipeline to find lobule-specific electric field distribution during non-invasive cerebellar stimulation. 2019 IEEE 16th International Conference on Rehabilitation Robotics (ICORR). 2019:1191–6.

[67] Bai S, Dokos S, Ho KA, Loo C. A computational modelling study of transcranial direct current stimulation montages used in depression. Neuroimage 2014;87:332–44.

[68] Rezaee Z, Dutta A. Cerebellar Lobules Optimal Stimulation (CLOS): A Computational Pipeline to Optimize Cerebellar Lobule-Specific Electric Field Distribution. Front Neurosci 2019;13.

[69] Bhalerao GV, Sreeraj VS, Bose A, Narayanaswamy JC, Venkatasubramanian G. Comparison of electric field modeling pipelines for transcranial direct current stimulation. Neurophysiol Clin 2021;51(4):303–18.

[70] Steinmann I, Williams KA, Wilke M, Antal A. Detection of Transcranial Alternating Current Stimulation Aftereffects Is Improved by Considering the Individual Electric Field Strength and Self-Rated Sleepiness. Front Neurosci 2022;16.

[71] Soleimani G, Towhidkhah F, Oghabian MA, Ekhtiari H. DLPFC stimulation alters large-scale brain networks connectivity during a drug cue reactivity task: A tDCS-fMRI study. Front Syst Neurosci 2022;16.

[72] Salehinejad MA, Ghayerin E, Nejati V, Yavari F, Nitsche MA. Domain-specific Involvement of the Right Posterior Parietal Cortex in Attention Network and Attentional Control of ADHD: A Randomized, Cross-over, Sham-controlled tDCS Study. Neuroscience 2020;444:149–59.

[73] Csifcsák G, Boayue NM, Puonti O, Thielscher A, Mittner M. Effects of transcranial direct current stimulation for treating depression: A modeling study. J Affect Disord 2018;234:164–73.

[74] Gomez-Tames J, Asai A, Mikkonen M, Laakso I, Tanaka S, Uehara S, et al. Group-level and functional-region analysis of electric-field shape during cerebellar transcranial direct current stimulation with different electrode montages. J Neural Eng 2019;16(3):036001.

[75] Boayue NM, Csifcsák G, Puonti O, Thielscher A, Mittner M. Head models of healthy and depressed adults for simulating the electric fields of non-invasive electric brain stimulation. F1000Res 2018;7:704.

[76] Zanto TP, Jones KT, Ostrand AE, Hsu W-Y, Campusano R, Gazzaley A. Individual differences in neuroanatomy and neurophysiology predict effects of transcranial alternating current stimulation. Brain Stimul 2021;14(5):1317–29.

[77] Evans C, Zich C, Lee JSA, Ward N, Bestmann S. Inter-individual variability in current direction for common tDCS montages. Neuroimage 2022;260:119501.

[78] Salvador R, Wenger C, Miranda PC. Investigating the cortical regions involved in MEP modulation in tDCS. Front Cell Neurosci 2015;9:405.

[79] Solanki D, Rezaee Z, Dutta A, Lahiri U. Investigating the feasibility of cerebellar transcranial direct current stimulation to facilitate post-stroke overground gait performance in chronic stroke: a partial least-squares regression approach. J Neuroeng Rehabil 2021;18(1):18.

[80] Kuhnke P, Beaupain MC, Cheung VKM, Weise K, Kiefer M, Hartwigsen G. Left posterior inferior parietal cortex causally supports the retrieval of action knowledge. Neuroimage 2020;219:117041.

[81] Rezaee Z, Dutta A. Lobule-Specific Dosage Considerations for Cerebellar Transcranial Direct Current Stimulation During Healthy Aging: A Computational Modeling Study Using Age-Specific Magnetic Resonance Imaging Templates. Neuromodulation 2020;23(3):341–65.

[82] Indahlastari A, Kasinadhuni AK, Saar C, Castellano K, Mousa B, Chauhan M, et al. Methods to Compare Predicted and Observed Phosphene Experience in tACS Subjects. Neural Plast 2018;2018:8525706.

[83] Cancelli A, Cottone C, Giordani A, Asta G, Lupoi D, Pizzella V, et al. MRI-Guided Regional Personalized Electrical Stimulation in Multisession and Home Treatments. Front Neurosci 2018;12:284.

[84] Splittgerber M, Borzikowsky C, Salvador R, Puonti O, Papadimitriou K, Merschformann C, et al. Multichannel anodal tDCS over the left dorsolateral prefrontal cortex in a paediatric population. Sci Rep 2021;11(1):21512.

[85] Abellaneda-Pérez K, Vaqué-Alcázar L, Perellón-Alfonso R, Solé-Padullés C, Bargalló N, Salvador R, et al. Multifocal Transcranial Direct Current Stimulation Modulates Resting-State Functional Connectivity in Older Adults Depending on the Induced Current Density. Front Aging Neurosci 2021;13:725013.

[86] Maran M, Numssen O, Hartwigsen G, Zaccarella E. Online neurostimulation of Broca’s area does not interfere with syntactic predictions: A combined TMS-EEG approach to basic linguistic combination. Front Psychol 2022;13.

[87] Lang S, Gan LS, McLennan C, Kirton A, Monchi O, Kelly JJP. Preoperative Transcranial Direct Current Stimulation in Glioma Patients: A Proof of Concept Pilot Study. Front Neurol 2020;11:593950.

[88] Gomez-Tames J, Asai A, Hirata A. Significant group-level hotspots found in deep brain regions during transcranial direct current stimulation (tDCS): A computational analysis of electric fields. Clin Neurophysiol 2020;131(3):755–65.

[89] Mezger E, Brunoni AR, Hasan A, Häckert J, Strube W, Keeser D, et al. tDCS for auditory verbal hallucinations in a case of schizophrenia and left frontal lesion: efield simulation and clinical results. Neurocase 2020;26(4):241–7.

[90] Fujimoto S, Tanaka S, Laakso I, Yamaguchi T, Kon N, Nakayama T, et al. The Effect of Dual-Hemisphere Transcranial Direct Current Stimulation Over the Parietal Operculum on Tactile Orientation Discrimination. Front Behav Neurosci 2017;11.

[91] Mikkonen M, Laakso I, Sumiya M, Koyama S, Hirata A, Tanaka S. TMS Motor Thresholds Correlate With TDCS Electric Field Strengths in Hand Motor Area. Front Neurosci 2018;12:426-.

[92] Zhang X, Hancock R, Santaniello S. Transcranial direct current stimulation of cerebellum alters spiking precision in cerebellar cortex: A modeling study of cellular responses. PLoS Comput Biol 2021;17(12):e1009609.

[93] Zmeykina E, Mittner M, Paulus W, Turi Z. Weak rTMS-induced electric fields produce neural entrainment in humans. Sci Rep 2020;10(1):11994.

[94] Bungert A, Antunes A, Espenhahn S, Thielscher A. Where does TMS Stimulate the Motor Cortex? Combining Electrophysiological Measurements and Realistic Field Estimates to Reveal the Affected Cortex Position. Cereb Cortex 2017;27(11):5083–94.

[95] Beynel L, Dannhauer M, Palmer H, Hilbig SA, Crowell CA, Wang JEH, et al. Network-based rTMS to modulate working memory: The difficult choice of effective parameters for online interventions. Brain and Behavior 2021;11(11):e2361.

[96] Nummenmaa A, Stenroos M, Ilmoniemi RJ, Okada YC, Hämäläinen MS, Raij T. Comparison of spherical and realistically shaped boundary element head models for transcranial magnetic stimulation navigation. Clin Neurophysiol 2013;124(10):1995–2007.

[97] van der Burght CL, Numssen O, Schlaak B, Goucha T, Hartwigsen G. Differential contributions of inferior frontal gyrus subregions to sentence processing guided by intonation. Hum Brain Mapp 2022;n/a(n/a).

[98] Passera B, Chauvin A, Raffin E, Bougerol T, David O, Harquel S. Exploring the spatial resolution of TMS-EEG coupling on the sensorimotor region. Neuroimage 2022;259:119419.

[99] Pizem D, Novakova L, Gajdos M, Rektorova I. Is the vertex a good control stimulation site? Theta burst stimulation in healthy controls. J Neural Transm 2022;129.

[100] Fiocchi S, Longhi M, Ravazzani P, Roth Y, Zangen A, Parazzini M. Modelling of the Electric Field Distribution in Deep Transcranial Magnetic Stimulation in the Adolescence, in the Adulthood, and in the Old Age. Comput Math Methods Med 2016;2016:9039613.

[101] Park J, Lee C, Lee S, Im C-H. 80 Hz but not 40 Hz, transcranial alternating current stimulation of 80 Hz over right intraparietal sulcus increases visuospatial working memory capacity. Sci Rep 2022;12(1):13762.

[102] Woods AJ, Bryant V, Sacchetti D, Gervits F, Hamilton R. Effects of Electrode Drift in Transcranial Direct Current Stimulation. Brain Stimul 2015;8(3):515–9.

[103] Benussi A, Cantoni V, Grassi M, Brechet L, Michel CM, Datta A, et al. Increasing Brain Gamma Activity Improves Episodic Memory and Restores Cholinergic Dysfunction in Alzheimer’s Disease. Ann Neurol 2022;92(2):322–34.

[104] Laakso I, Tanaka S, Koyama S, De Santis V, Hirata A. Inter-subject Variability in Electric Fields of Motor Cortical tDCS. Brain Stimul 2015;8(5):906–13.

[105] Stoupis D, Samaras T. Non-invasive stimulation with temporal interference: optimization of the electric field deep in the brain with the use of a genetic algorithm. J Neural Eng 2022;19(5):056018.

[106] Gillick BT, Kirton A, Carmel JB, Minhas P, Bikson M. Pediatric stroke and transcranial direct current stimulation: methods for rational individualized dose optimization. Front Hum Neurosci 2014;8:739.

[107] Edwards D, Cortes M, Datta A, Minhas P, Wassermann EM, Bikson M. Physiological and modeling evidence for focal transcranial electrical brain stimulation in humans: a basis for high-definition tDCS. Neuroimage 2013;74:266–75.

[108] Seo H, Jun SC. Relation between the electric field and activation of cortical neurons in transcranial electrical stimulation. Brain Stimul 2019;12(2):275–89.

[109] DaSilva AF, Truong DQ, DosSantos MF, Toback RL, Datta A, Bikson M. State-of-art neuroanatomical target analysis of high-definition and conventional tDCS montages used for migraine and pain control. Front Neuroanat 2015;9:89.

[110] Manoli Z, Parazzini M, Ravazzani P, Samaras T. The electric field distributions in anatomical head models during transcranial direct current stimulation for post-stroke rehabilitation. Med Phys 2017;44(1):262–71.

[111] Seibt O, Brunoni AR, Huang Y, Bikson M. The Pursuit of DLPFC: Non-neuronavigated Methods to Target the Left Dorsolateral Pre-frontal Cortex With Symmetric Bicephalic Transcranial Direct Current Stimulation (tDCS). Brain Stimul 2015;8(3):590–602.

[112] da Silva RMF, Batistuzzo MC, Shavitt RG, Miguel EC, Stern E, Mezger E, et al. Transcranial direct current stimulation in obsessive-compulsive disorder: an update in electric field modeling and investigations for optimal electrode montage. Expert Rev Neurother 2019;19(10):1025–35.

[113] Kalloch B, Bazin P-L, Villringer A, Sehm B, Hlawitschka M. A flexible workflow for simulating transcranial electric stimulation in healthy and lesioned brains. PLoS One 2020;15(5):e0228119.

[114] Colella M, Paffi A, De Santis V, Apollonio F, Liberti M. Effect of skin conductivity on the electric field induced by transcranial stimulation techniques in different head models. Physics in Medicine & Biology 2021;66(3):035010.

[115] Jamil A, Batsikadze G, Kuo H-I, Meesen RLJ, Dechent P, Paulus W, et al. Current intensity-and polarity-specific online and aftereffects of transcranial direct current stimulation: An fMRI study. Hum Brain Mapp 2020;41(6):1644–66.

[116] Beynel L, Davis SW, Crowell CA, Dannhauer M, Lim W, Palmer H, et al. Site-Specific Effects of Online rTMS during a Working Memory Task in Healthy Older Adults. Brain Sci 2020;10(5).

[117] Lee EG, Rastogi P, Hadimani RL, Jiles DC, Camprodon JA. Impact of non-brain anatomy and coil orientation on inter-and intra-subject variability in TMS at midline. Clin Neurophysiol 2018;129(9):1873–83.

[118] Preisig B, Hervais-Adelman A. The Predictive Value of Individual Electric Field Modeling for Transcranial Alternating Current Stimulation Induced Brain Modulation. Front Cell Neurosci 2022;16.

[119] Alekseichuk I, Mantell K, Shirinpour S, Opitz A. Comparative modeling of transcranial magnetic and electric stimulation in mouse, monkey, and human. Neuroimage 2019;194:136–48.

[120] Li H, Deng Z-D, Oathes D, Fan Y. Computation of transcranial magnetic stimulation electric fields using self-supervised deep learning. Neuroimage 2022;264:119705.

[121] Uenishi S, Tamaki A, Yamada S, Yasuda K, Ikeda N, Mizutani-Tiebel Y, et al. Computational modeling of electric fields for prefrontal tDCS across patients with schizophrenia and mood disorders. Psychiatry Res Neuroimaging 2022;326:111547.

[122] Wysokiński A. Does sponge pads wear affect the distribution of electric field generated by tDCS? Clin Neurophysiol 2021;132(8):1782–4.

[123] Antonenko D, Thams F, Grittner U, Uhrich J, Glöckner F, Li SC, et al. Randomized trial of cognitive training and brain stimulation in non-demented older adults. Alzheimers Dement (N Y) 2022;8(1):e12262.

[124] Miranda PC, Mekonnen A, Salvador R, Ruffini G. The electric field in the cortex during transcranial current stimulation. Neuroimage 2013;70:48–58.

[125] Scheldrup M, Greenwood PM, McKendrick R, Strohl J, Bikson M, Alam M, et al. Transcranial direct current stimulation facilitates cognitive multi-task performance differentially depending on anode location and subtask. Front Hum Neurosci 2014;8:665.

[126] Lee WH, Kennedy NI, Bikson M, Frangou S. A Computational Assessment of Target Engagement in the Treatment of Auditory Hallucinations with Transcranial Direct Current Stimulation. Front Psychiatry 2018;9:48.

[127] Carla Piastra M, van der Cruijsen J, Piai V, Jeukens FEM, Manoochehri M, Schouten AC, et al. ASH: an Automatic pipeline to generate realistic and individualized chronic Stroke volume conduction Head models. J Neural Eng 2021;18(4).

[128] Shahid S, Wen P, Ahfock T. Assessment of electric field distribution in anisotropic cortical and subcortical regions under the influence of tDCS. Bioelectromagnetics 2014;35(1):41–57.

[129] Soldati M, Laakso I. Computational errors of the induced electric field in voxelized and tetrahedral anatomical head models exposed to spatially uniform and localized magnetic fields. Phys Med Biol 2020;65(1):015001.

[130] Tzirini M, Roth Y, Harmelech T, Zibman S, Pell GS, Kimiskidis VK, et al. Detailed measurements and simulations of electric field distribution of two TMS coils cleared for obsessive compulsive disorder in the brain and in specific regions associated with OCD. PLoS One 2022;17(8):e0263145.

[131] Mizutani-Tiebel Y, Takahashi S, Karali T, Mezger E, Bulubas L, Papazova I, et al. Differences in electric field strength between clinical and non-clinical populations induced by prefrontal tDCS: A cross-diagnostic, individual MRI-based modeling study. Neuroimage Clin 2022;34:103011.

[132] Soleimani G, Saviz M, Bikson M, Towhidkhah F, Kuplicki R, Paulus MP, et al. Group and individual level variations between symmetric and asymmetric DLPFC montages for tDCS over large scale brain network nodes. Sci Rep 2021;11(1):1271.

[133] Antonenko D, Grittner U, Saturnino G, Nierhaus T, Thielscher A, Flöel A. Inter-individual and age-dependent variability in simulated electric fields induced by conventional transcranial electrical stimulation. Neuroimage 2021;224:117413.

[134] Alawi M, Lee PF, Deng Z-D, Goh YK, Croarkin PE. Modelling on differential effect of age on transcranial magnetic stimulation induced electric fields. J Neural Eng 2022.

[135] Parazzini M, Rossi E, Ferrucci R, Liorni I, Priori A, Ravazzani P. Modelling the electric field and the current density generated by cerebellar transcranial DC stimulation in humans. Clin Neurophysiol 2014;125(3):577–84.

[136] Klaus J, Schutter D. Putting focus on transcranial direct current stimulation in language production studies. PLoS One 2018;13(8):e0202730.

[137] Wang B, Shen MR, Deng ZD, Smith JE, Tharayil JJ, Gurrey CJ, et al. Redesigning existing transcranial magnetic stimulation coils to reduce energy: application to low field magnetic stimulation. J Neural Eng 2018;15(3):036022.

[138] Alam M, Truong DQ, Khadka N, Bikson M. Spatial and polarity precision of concentric high-definition transcranial direct current stimulation (HD-tDCS). Phys Med Biol 2016;61(12):4506–21.

[139] Metwally MK, Han SM, Kim TS. The effect of tissue anisotropy on the radial and tangential components of the electric field in transcranial direct current stimulation. Med Biol Eng Comput 2015;53(10):1085–101.

[140] Janssen AM, Rampersad SM, Lucka F, Lanfer B, Lew S, Aydin U, et al. The influence of sulcus width on simulated electric fields induced by transcranial magnetic stimulation. Phys Med Biol 2013;58(14):4881–96.

[141] Glasser MF, Coalson TS, Robinson EC, Hacker CD, Harwell J, Yacoub E, et al. A multi-modal parcellation of human cerebral cortex. Nature 2016;536(7615):171–8.

[142] McGrath H, Zaveri HP, Collins E, Jafar T, Chishti O, Obaid S, et al. High-resolution cortical parcellation based on conserved brain landmarks for localization of multimodal data to the nearest centimeter. Sci Rep 2022;12(1):18778.

[143] Josse G, Kherif F, Flandin G, Seghier ML, Price CJ. Predicting language lateralization from gray matter. J Neurosci 2009;29(43):13516–23.

[144] Cox SR, Ritchie SJ, Tucker-Drob EM, Liewald DC, Hagenaars SP, Davies G, et al. Ageing and brain white matter structure in 3,513 UK Biobank participants. Nat Commun 2016;7(1):13629.

[145] Indahlastari A, Albizu A, O’Shea A, Forbes MA, Nissim NR, Kraft JN, et al. Modeling transcranial electrical stimulation in the aging brain. Brain Stimul 2020;13(3):664–74.

[146] Ghasemian-Shirvan E, Farnad L, Mosayebi-Samani M, Verstraelen S, Meesen RLJ, Kuo MF, et al. Age-related differences of motor cortex plasticity in adults: A transcranial direct current stimulation study. Brain Stimul 2020;13(6):1588–99.

[147] Zatorre RJ, Fields RD, Johansen-Berg H. Plasticity in gray and white: neuroimaging changes in brain structure during learning. Nat Neurosci 2012;15(4):528–36.

[148] Sampaio-Baptista C, Khrapitchev AA, Foxley S, Schlagheck T, Scholz J, Jbabdi S, et al. Motor skill learning induces changes in white matter microstructure and myelination. J Neurosci 2013;33(50):19499–503.

