## Supplementary materials for "A Systematic Review and Large-Scale tES and TMS Electric Field Modeling Study Reveals How Outcome Measure Selection Alters Results in a Person- and Montage-Specific Manner"

**Supplementary materials 1.** **Search keys**

PubMed

((((((electrical field[Title/Abstract]) OR (electrical fields[Title/Abstract])) OR (E-field[Title/Abstract])) OR (E-fields[Title/Abstract])) OR (electric field[Title/Abstract])) OR (electric fields[Title/Abstract])) AND (((((((((((noninvasive brain stimulation[Title/Abstract]) OR (non-invasive brain stimulation[Title/Abstract])) OR (non invasive brain stimulation[Title/Abstract])) OR (NIBS[Title/Abstract])) OR (tDCS[Title/Abstract])) OR (TMS[Title/Abstract])) OR (tACS[Title/Abstract])) OR (tES[Title/Abstract])) OR (transcranial magnetic stimulation[Title/Abstract])) OR (transcranial direct current stimulation[Title/Abstract])) OR (transcranial alternating current stimulation[Title/Abstract]))

Scopus

( ( TITLE-ABS-KEY ( noninvasive AND brain AND stimulation ) OR TITLE-ABS-KEY ( nibs ) OR TITLE-ABS-KEY ( non-invasive AND brain AND stimulation ) OR TITLE-ABS-KEY ( tdcs ) OR TITLE-ABS-KEY ( tms ) OR TITLE-ABS-KEY ( tacs ) OR TITLE-ABS-KEY ( tes ) OR TITLE-ABS-KEY ( transcranial AND magnetic AND stimulation ) OR TITLE-ABS-KEY ( transcranial AND direct AND current AND stimulation ) OR TITLE-ABS-KEY ( transcranial AND alternating AND current AND stimulation ) OR TITLE-ABS-KEY ( non AND invasive AND brain AND stimulation ) ) AND PUBYEAR > 2011 AND PUBYEAR > 2011 ) AND ( ( TITLE-ABS-KEY ( electrical AND field ) OR TITLE-ABS-KEY ( electrical AND fields ) OR TITLE-ABS-KEY ( e-fields ) OR TITLE-ABS-KEY ( e-field ) OR TITLE-ABS-KEY ( electric AND field ) OR TITLE-ABS-KEY ( electric AND fields ) ) AND PUBYEAR > 2011 AND PUBYEAR > 2011 ) AND ( LIMIT-TO ( DOCTYPE , "ar" ) OR LIMIT-TO ( DOCTYPE , "cp" ) OR LIMIT-TO ( DOCTYPE , "re" ) OR LIMIT-TO ( DOCTYPE , "le" ) ) AND ( LIMIT-TO ( SUBJAREA , "NEUR" ) ) AND ( LIMIT-TO ( LANGUAGE , "English" ) )

Web Of Science

((TS=(electric field) OR TS=(electric fields) OR TS=(E-field) OR TS=(E-FIELDS) OR TS=(electric fields)) AND (TS=(noninvasive brain stimulation) OR TS=(non-invasive brain stimulation) OR TS=(non invasive brain stimulation) OR TS=(NIBS) OR TS=(tDCS) OR TS=(TMS) OR TS=(tACS) OR TS=(tES) OR TS=(transcranial magnetic stimulation) OR TS=(transcranial direct current stimulation) OR TS=(transcranial alternating current stimulation)) AND (PY==("2022" OR "2021" OR "2020" OR "2019" OR "2018" OR "2017" OR "2016" OR "2015" OR "2014" OR "2013" OR "2012") AND TASCA==("CLINICAL NEUROLOGY" OR "NEUROSCIENCES") AND DT==("REVIEW" OR "ARTICLE") AND LA==("ENGLISH"))

**Supplementary materials 2. Conductivity values used for electric field simulations**

| **Tissue Name** | **Value (S/m)** | **Tissue Name** | **Value (S/m)** |
| --- | --- | --- | --- |
| White Matter | 0.126 | Compact Bone | 0.008 |
| Gray Matter | 0.275 | Spongy Bone | 0.025 |
| CSF | 1.654 | Blood | 0.600 |
| Bone | 0.010 | Muscle | 0.160 |
| Scalp | 0.465 | Silicone Rubber | 29.400 |
| Eye balls | 0.500 | Saline | 1.000 |

**Supplementary materials 3. Identification of peak electric field value (V/m)**

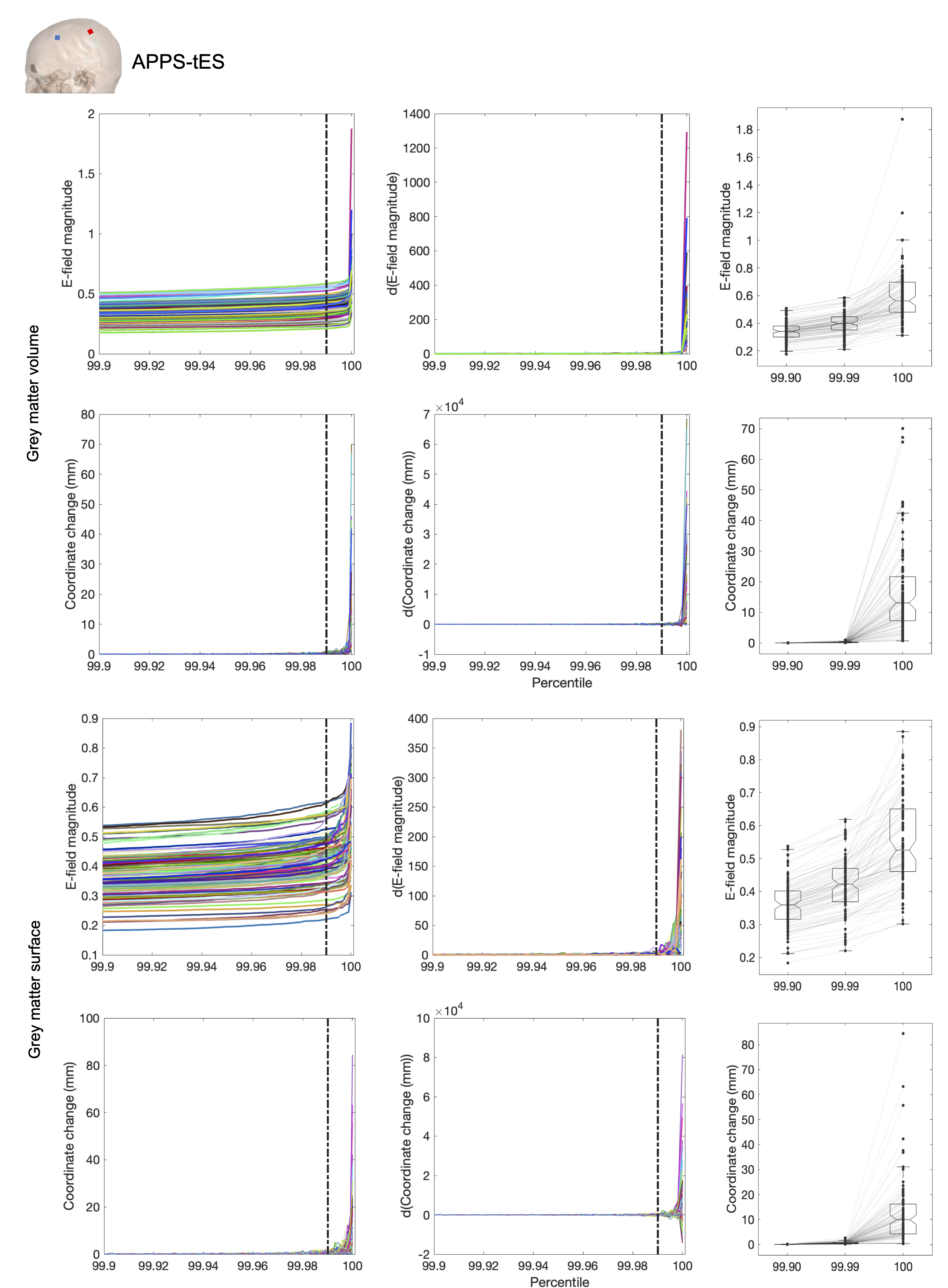

**Figure S3.1.** Evaluation of different electric field magnitude values and the associated 3D location across different percentiles used to approximate the peak electric field magnitude. P99.9 was withheld as peak electric field value due to it being the highest value that was stable both across the grey matter volume and surface, that was also used in several previous studies.

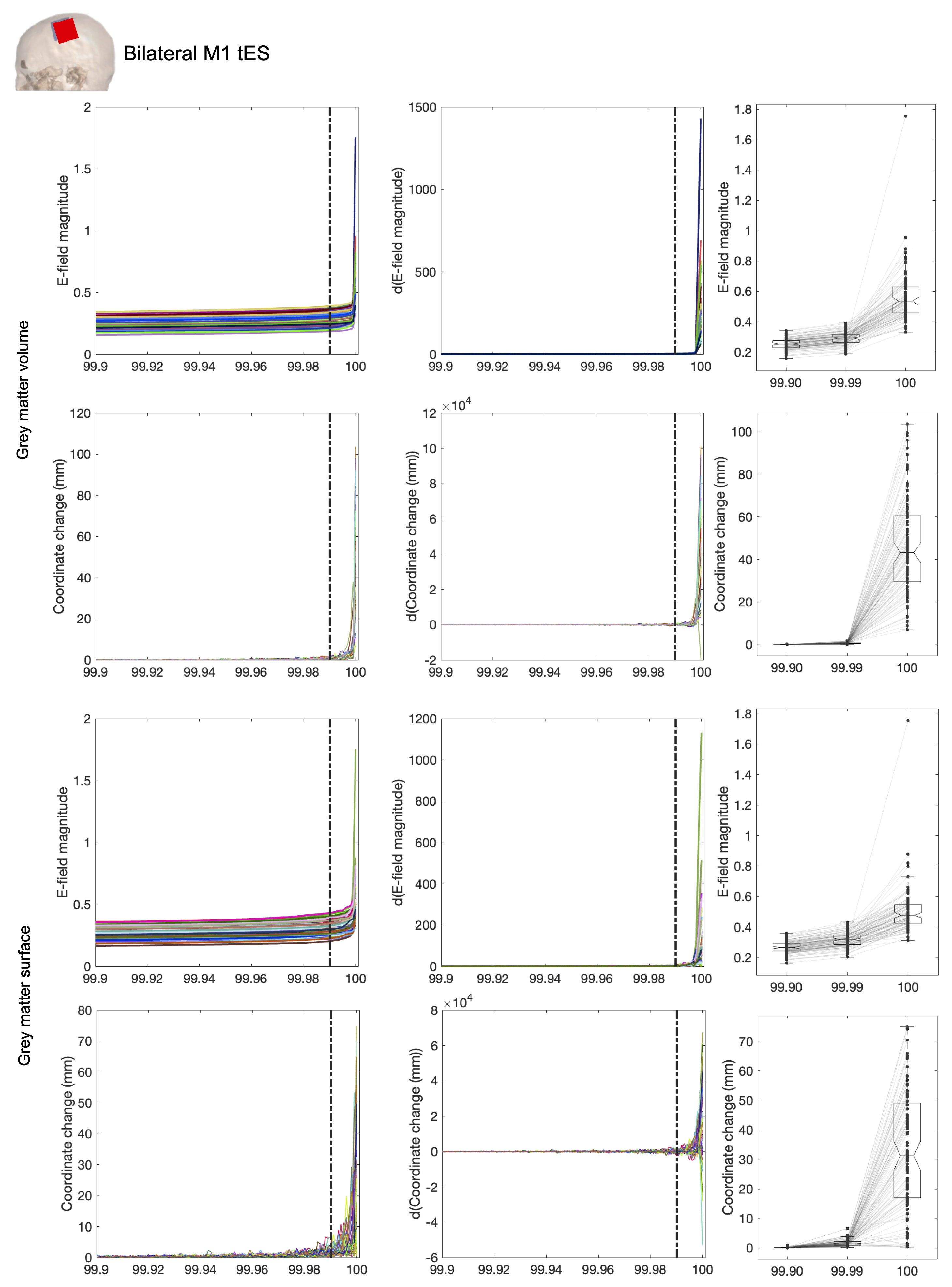

**Figure S3.2.** Evaluation of different electric field magnitude values and the associated 3D location across different percentiles used to approximate the peak electric field magnitude. P99.9 was withheld as peak electric field value due to it being the highest value that was stable both across the grey matter volume and surface, that was also used in several previous studies.

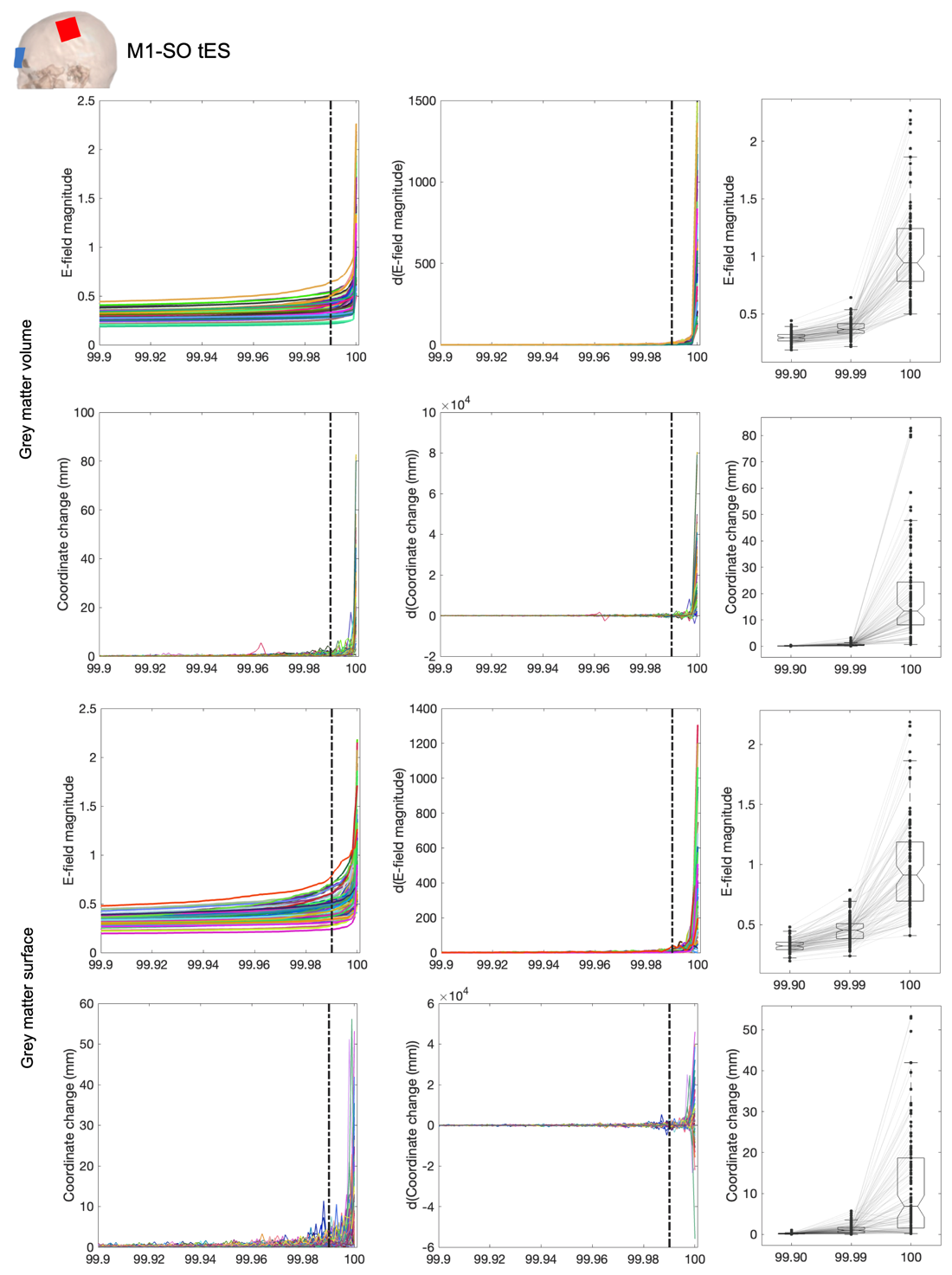

**Figure S3.3.** Evaluation of different electric field magnitude values and the associated 3D location across different percentiles used to approximate the peak electric field magnitude. P99.9 was withheld as peak electric field value due to it being the highest value that was stable both across the grey matter volume and surface, that was also used in several previous studies.

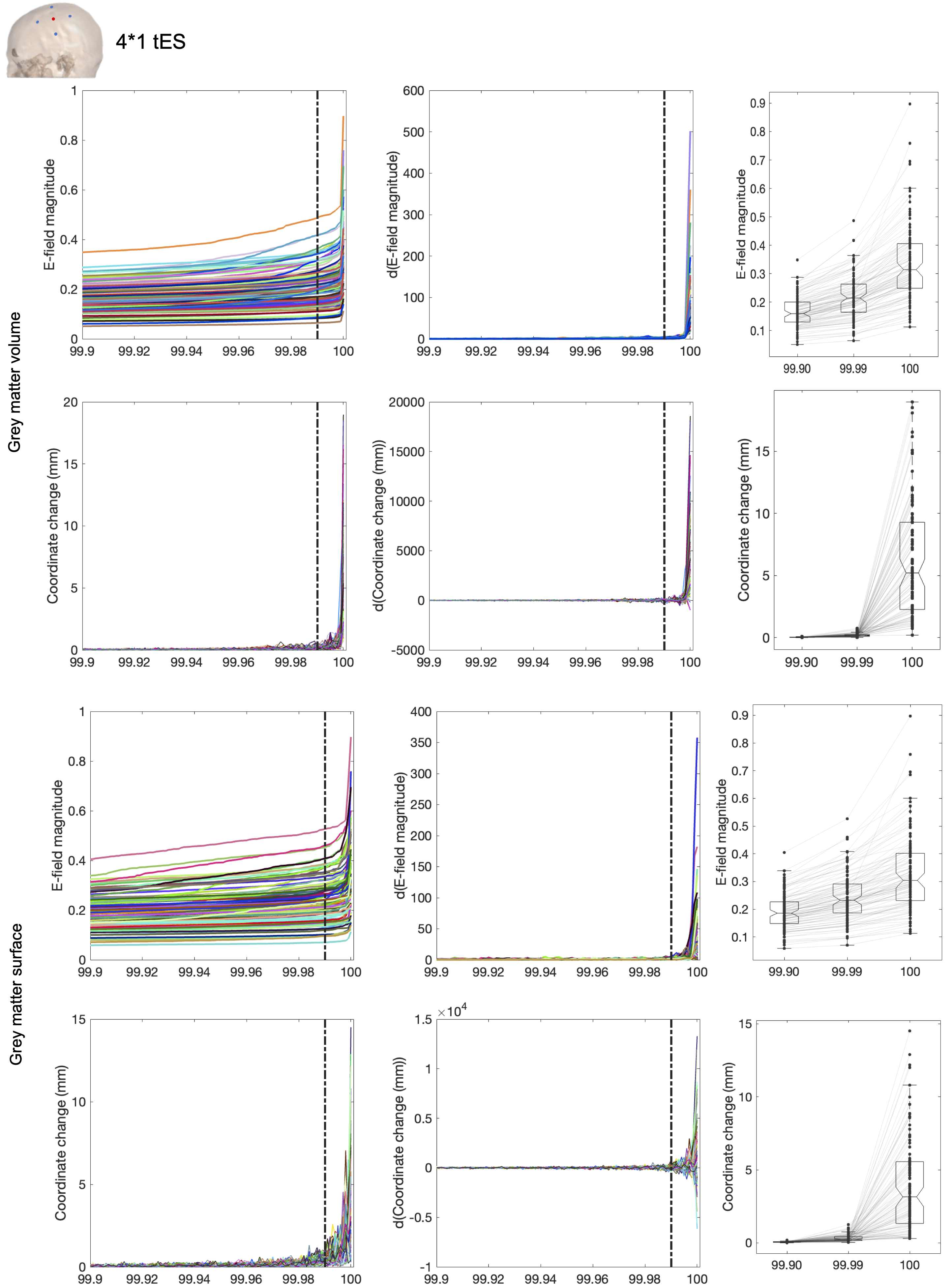

**Figure S3.4.** Evaluation of different electric field magnitude values and the associated 3D location across different percentiles used to approximate the peak electric field magnitude. P99.9 was withheld as peak electric field value due to it being the highest value that was stable both across the grey matter volume and surface, that was also used in several previous studies.

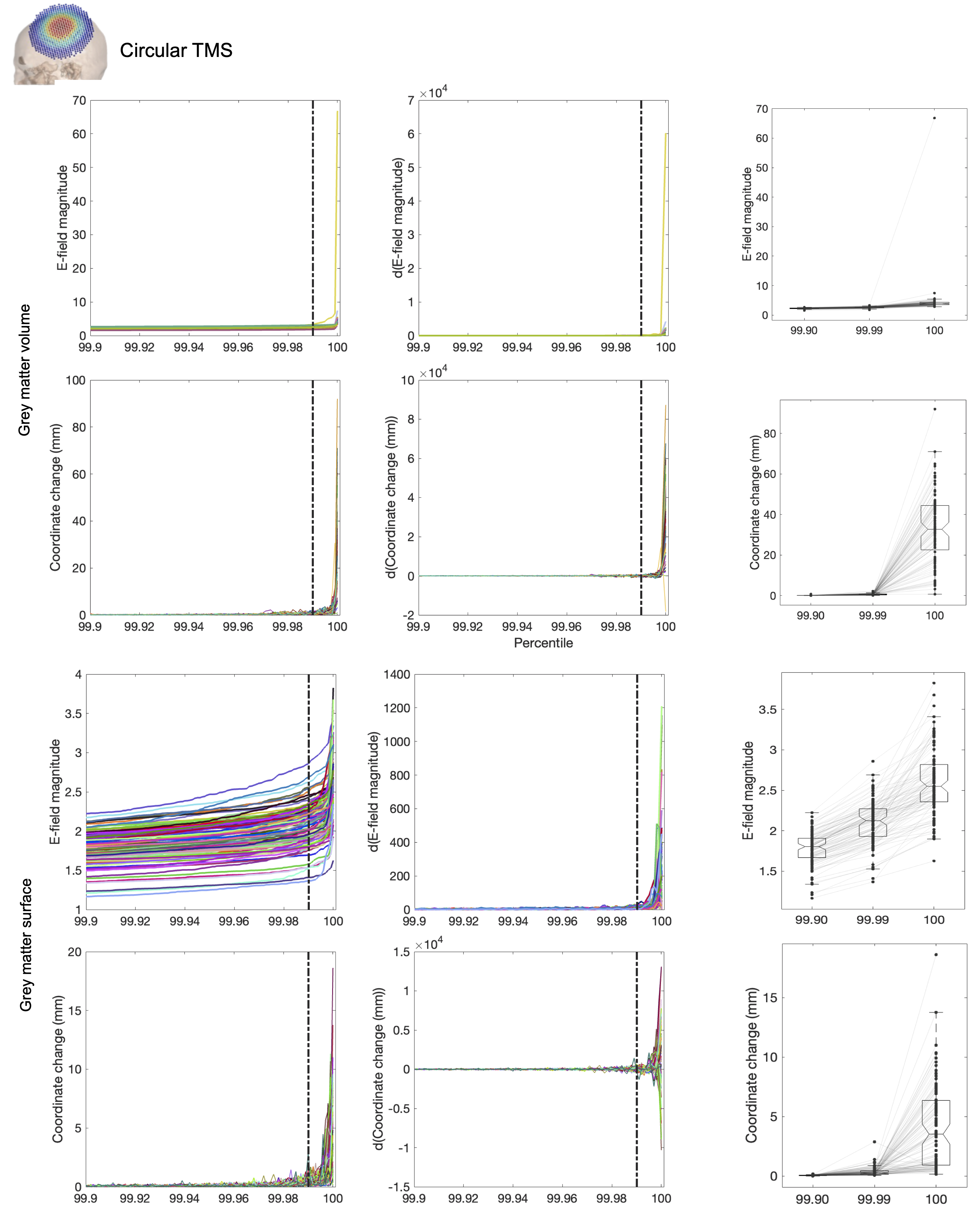

**Figure S3.5.** Evaluation of different electric field magnitude values and the associated 3D location across different percentiles used to approximate the peak electric field magnitude. P99.9 was withheld as peak electric field value due to it being the highest value that was stable both across the grey matter volume and surface, that was also used in several previous studies. Here, we show the results at a stimulation intensity of 1 dI/dt. Thus, these results can be easily scaled by multiplying with the wanted intensity.

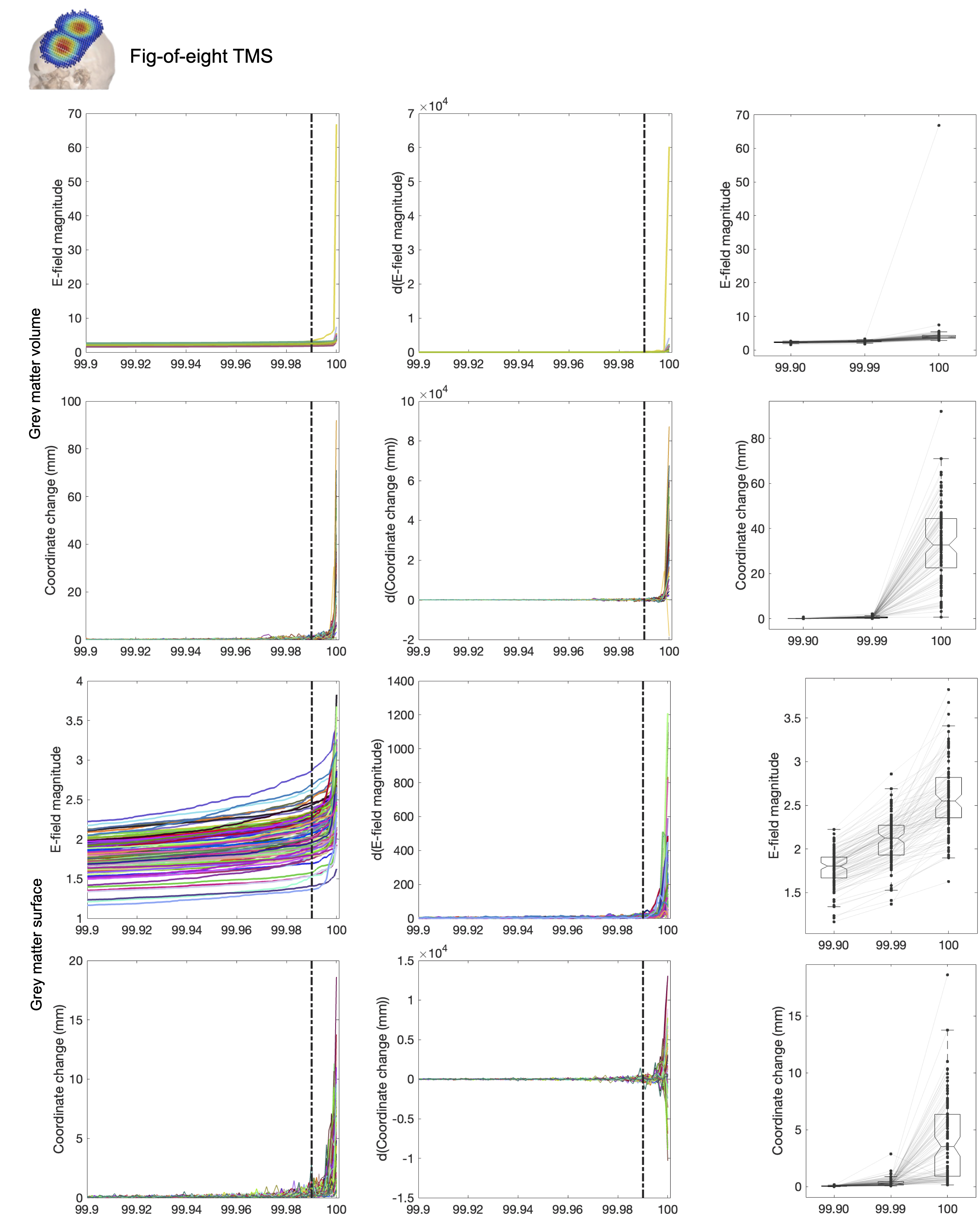

**Figure S3.6.** Evaluation of different electric field magnitude values and the associated 3D location across different percentiles used to approximate the peak electric field magnitude. P99.9 was withheld as peak electric field value due to it being the highest value that was stable both across the grey matter volume and surface, that was also used in several previous studies. Here, we show the results at a stimulation intensity of 1 dI/dt. Thus, these results can be easily scaled by multiplying with the wanted intensity.
